# The laminar profile of sleep spindles in humans

**DOI:** 10.1101/563221

**Authors:** Péter P. Ujma, Boglárka Hajnal, Róbert Bódizs, Ferenc Gombos, Loránd Erőss, Lucia Wittner, Eric Halgren, Sydney Cash, István Ulbert, Dániel Fabó

**Affiliations:** Institute of Behavioural Sciences, Semmelweis University, Budapest, Hungary; “Juhász Pál” Epilepsy Centrum, National Institute of Clinical Neurosciences, Budapest, Hungary; School of P.h.D. studies, Semmelweis University, Budapest, Hungary; Institute of Cognitive Neuroscience and Psychology, Research Center for Natural Sciences, Hungarian Academy of Sciences, Budapest, Hungary; Departments of Radiology and Neurosciences, University of California, San Diego CA, USA; Center for Neurotechnology and Neurorecovery (CNTR), Department of Neurology, Massachusetts General Hospital, Boston, MA, USA; Department of Neurology, Harvard Medical School, Boston, MA, USA

## Abstract

Sleep spindles are generated in thalamocortical, corticothalamic and possibly cortico-cortical circuits. Previous hypotheses suggested that slow and fast spindles or spindles with various spatial extent may be generated in different circuits with various cortical laminar innervation patterns. We used NREM sleep EEG data recorded from four human epileptic patients undergoing presurgical electrophysiological monitoring with subdural electrocorticographic grids (ECoG) and implanted laminar microelectrodes penetrating the cortex (IME). The position of IME electrodes within cortical layers was confirmed using postsurgical histological reconstructions. Many micro-domain spindles detected on the IME occurred only in one layer and were absent from the ECoG, but with increasing amplitude simultaneous detection in other layers and on the ECoG became more likely. Macro-domain spindles sufficiently large to be detected on the ECoG were in contrast usually accompanied by IME spindles. Neither micro-domain nor macro-domain spindle cortical profiles were strongly associated with sleep spindle frequency or globality. Multiple-unit and single-unit activity during spindles, however, was heterogeneous across spindle types, but also across layers and subjects. Our results indicate that extremely local spindles may occur in any cortical layer, but co-occurrence at other locations becomes likelier with increasing amplitude and the relatively large spindles detected on ECoG channels have a stereotypical laminar profile. We found no compelling evidence that different spindle types are associated with different laminar profiles, suggesting that they are generated in cortical and thalamic circuits with similar cortical innervation patterns. Local neuronal activity is a stronger candidate mechanism for driving functional differences between spindles subtypes.

## Introduction

Sleep spindles are roughly sinusoid oscillations arising from the interaction of thalamocortical, corticothalamic and reticular thalamic networks (Steriade, 2003; Fogel and Smith, 2011; Lüthi, 2013), visible on both scalp EEG and ECoG recordings (Nakamura et al., 2003; Andrillon et al., 2011; Fogel and Smith, 2011; Peter-Derex et al., 2012; Frauscher et al., 2015). Sleep spindles are characterized by neural firing patterns which are highly conductive for long-term synaptic changes (Lüthi, 2013), including long-term potentiation (LTP) (Rosanova and Ulrich, 2005), which is usually suppressed in NREM sleep as a result of decreased cholinergic activation (Tononi and Cirelli, 2014). Sleep spindles have been extensively studied for their role in memory, cognition and sleep function. Sleep spindles have been implicated in learning and its efficacy (Gais et al., 2002; Gais and Born, 2004; Clemens et al., 2005), play a role in general cognitive ability (Bodizs et al., 2005; Schabus et al., 2006; Bόdizs et al., 2014; Ujma et al., 2014; Ujma et al., 2015c), but they are also important clinical markers in both neurological and psychiatric conditions (Bόdizs et al., 2012; Ferrarelli, 2015; Manoach et al., 2015; Gorgoni et al., 2016; Berencsi et al., 2017).

As sleep spindles arise in thalamocortical networks, they are subject to the anatomical properties of thalamocortical connections. Based on the existence of ‘core’ and ‘matrix’ thalamocortical connections (Jones, 1998, 2001) - which project to cortical layer IV and I-II, respectively - it has been hypothesized that sleep spindles can arise through either or both of these networks, possibly with a highly variable contribution of each network across spindles (Piantoni et al., 2016). Since different thalamocortical projections terminate in different cortical layers, the thalamic source of an electroencephalographic event can be inferred from sinks and sources in the current source density of signals recorded from electrodes which penetrate the cerebral cortex (Freeman and Nicholson, 1975; Ulbert et al., 2001). Studies in human epileptic patients have successfully revealed the laminar profile of the K-complex (Cash et al., 2009) and the slow wave (Csercsa et al., 2010), both of which were revealed to originate mainly from superficial layers. Results from animal laminar recordings (Spencer and Brookhart, 1961; Kandel and Buzsaki, 1997) - while not explicitly investigating this in the framework of the core-matrix dichotomy - reported that various thalamocortical networks are involved in sleep spindle generation, evidenced by a main sink during spindles in layer IV together with a more superficial sink. Recently, a study of human epileptic subjects with laminar probes (Hagler et al., 2018) found that sleep spindles recorded from the cortex occur with variable topographies and some spindles are localized to either upper or middle layers of the cortex.

Sleep spindles can be divided into slow and fast subtypes. Slow and fast spindles are clearly separated by frequency and topography (Andrillon et al., 2011; Fogel and Smith, 2011), characterized by different cerebral hemodynamic responses (Schabus et al., 2007), and coupled differently to the cortical slow oscillation (Mölle et al., 2011). There is substantial variability across individuals in the frequency of sleep spindles, especially slow spindles (De Gennaro et al., 2008; Ujma et al., 2015a). In spite of this fact, not all studies analyze slow and fast spindles separately, and only very few take into account the individual variability in sleep spindle frequency. Furthermore, there is substantial variability in the spatial extent of spindles (Andrillon et al., 2011; Nir et al., 2011; Piantoni et al., 2017).

Some limited evidence is available on how the thalamocortical generators of spindles diverge in slow and fast spindles or spindle with a different degree of cortical involvement. First, an in silico simulation of thalamocortical networks (Bonjean et al., 2012) demonstrated that more widespread thalamocortical connections to superficial layers contribute to the greater spatial extent and cortical synchrony of spindles. Second, pharmacological manipulations can impact affect slow and fast spindles differently: the Ca2+ channel antagonist flunarazine selectively reduces fast spindle activity, while the voltage-gated Na+ channel antagonist carbamazepine not only reduces fast spindle activity, but actually enhances both slow spindle and ∼0.75 Hz slow wave activity (Ayoub et al., 2013). Notably, Ca2+ channels are implicated in the *thalamic* generation of spindles (Astori and Luthi, 2013; Lüthi, 2013), especially in the reticular nucleus (TNR): thus, the resilience of slow spindles to the blocking of these channels suggests that they can be generated independently of the thalamus, in substantial contrast to traditional theories of spindle generation (McCormick and Bal, 1994; Steriade, 2003). This has led to the hypothesis (Timofeev and Chauvette, 2013) that slow spindle generators differ from fast spindle generators and possibly overlap with slow wave generators, which project to superficial cortical layers (Csercsa et al., 2010).

Therefore, it can be hypothesized that slow spindles and widespread (global) spindles have greater reliance on thalamocortical and possibly cortico-cortical networks terminating in superficial layers (“matrix” network), while fast spindles and local spindles preferentially rely on the more spatially focused “core” network (Piantoni et al., 2016). In line with this theory, relatively more superficial activations would be expected in case of slow and global spindles, while the activation should be higher in deeper cortical layers during fast and local spindles. It has recently been demonstrated that spindle-frequency oscillations recorded from the depth of the cortex indeed occur with a heterogeneous topography (Hagler et al., 2018). However, it is unknown whether different sleep spindle topographies are correlated with spindle subtypes.

The aim of our study with simultaneous electrocorticographic (ECoG) and laminar (IME) recordings in humans was to determine how neural networks with afferentation to different layers contribute to sleep spindle generation, and whether this contribution is different in different spindle types (slow/fast, local/global). We performed analyses to determine the dependence of the cortical profile of both “micro-domain” (IME) and “macro-domain” (ECoG) spindles on sleep spindle type.

## Methods

### Patients and data selection

Sleep electrophysiological data from four patients with drug-resistant epilepsy undergoing presurgical electrophysiological monitoring was selected for analysis. The selection criteria for patients were the following: 1) simultaneous recordings of electrocorticography (ECoG) data from subdural grids and at least one implanted microelectrode (IME) 2) existing post-surgical histological reconstruction of the neural tissue surrounding the IME (electrode track). The selection criteria for EEG data were the following: 1) seizure-free data with clearly identifiable NREM sleep, defined by the presence of sleep spindles and slow waves on ECoG and (if available) scalp EEG channels 2) adequate signal quality of both ECoG and IME data, indicated by the absence of continuous, broad-frequency artifacts. Furthermore, the absence of high-frequency artifacts (verified by visual inspection) and the presence of visible single-unit peaks were the prerequisite for considering the data from a subject for multiple unit and single-unit activity analysis, respectively. For the simultaneous analysis of ECoG EEG and IME MUA data in Patient 1-2, only ECoG spindles concomitant to high-quality IME signal was used. Postoperative examinations confirmed that the IME was implanted outside the seizure onset zone in Patients 1-3, but within the seizure onset zone in Patient 4.

All interventions were approved by the Hungarian Medical Scientific Council and the ethical committee of the National Institute of Clinical Neuroscience. Clinical procedures were not biased for scientific purposes. All patients gave informed consent in line with the Declaration of Helsinki.

### Implantation anatomy and histology

Electrode positions were confirmed by intraoperative navigation, the comparison of pre-and postoperative MR scans, as well as the comparison of photographs taken during the initial surgery and the ones taken during resective surgery. The brain tissue containing the IME was removed during surgery, cut to 2-5 mm blocks and chemically fixated (Ulbert et al., 2004; Csercsa et al., 2010). The laminar topography of the cortical tissue blocks was reconstructed from these samples in all patients and it was used to determine the location of IME electrodes within cortical layers, taking into account the shrinkage of brain tissue during preparation (Wittner et al., 2006; Csercsa et al., 2010). The cortical layer from which an IME signal originated was determined based on the histological reconstruction of the electrode track.

### Electrophysiology

All patients underwent electrophysiological recordings using implanted laminar microelectrodes (IME, 24 electrodes) and subdural grid and strip electrodes, from which only grids were analyzed (ECoG, 20-64 electrodes) **(Figure 1).** IMEs had a diameter of 350 μm, inserted perpendicularly to the cortical surface, penetrating the cortex to a depth of 3.5 mm. 40 μm platinum/iridium electrodes were built into the IME, spaced evenly at 150 μm (Ulbert et al., 2001). A silicone sheet at the top of the microelectrode prevented the array from shifting below the pial surface. ECoG hardware filters were set to 0.1-200 Hz (ECoG). ECoG data was recorded with a contralateral mastoid reference. IME data was recorded with a bipolar reference, with each contact referred to the one inferior to it (local field potential gradient, LFPg), yielding an effectively reference-free recording (Ulbert et al., 2001). IME data was separated into the EEG range (0.1-300 Hz) and the single/multiple unit range (300-5000 Hz) at the level of the amplifier and recorded as two separate data files (Ulbert et al., 2001; Cash et al., 2009; Keller et al., 2009; Csercsa et al., 2010). ECoG was recorded with a sampling frequency of either 2000 Hz/16 bit (patient 1 and 2) or 1024 Hz/16 bit (patient 3 and 4), while IME data was recorded with a sampling frequency of 20 kHz/12 bit.

**Figure 1.**
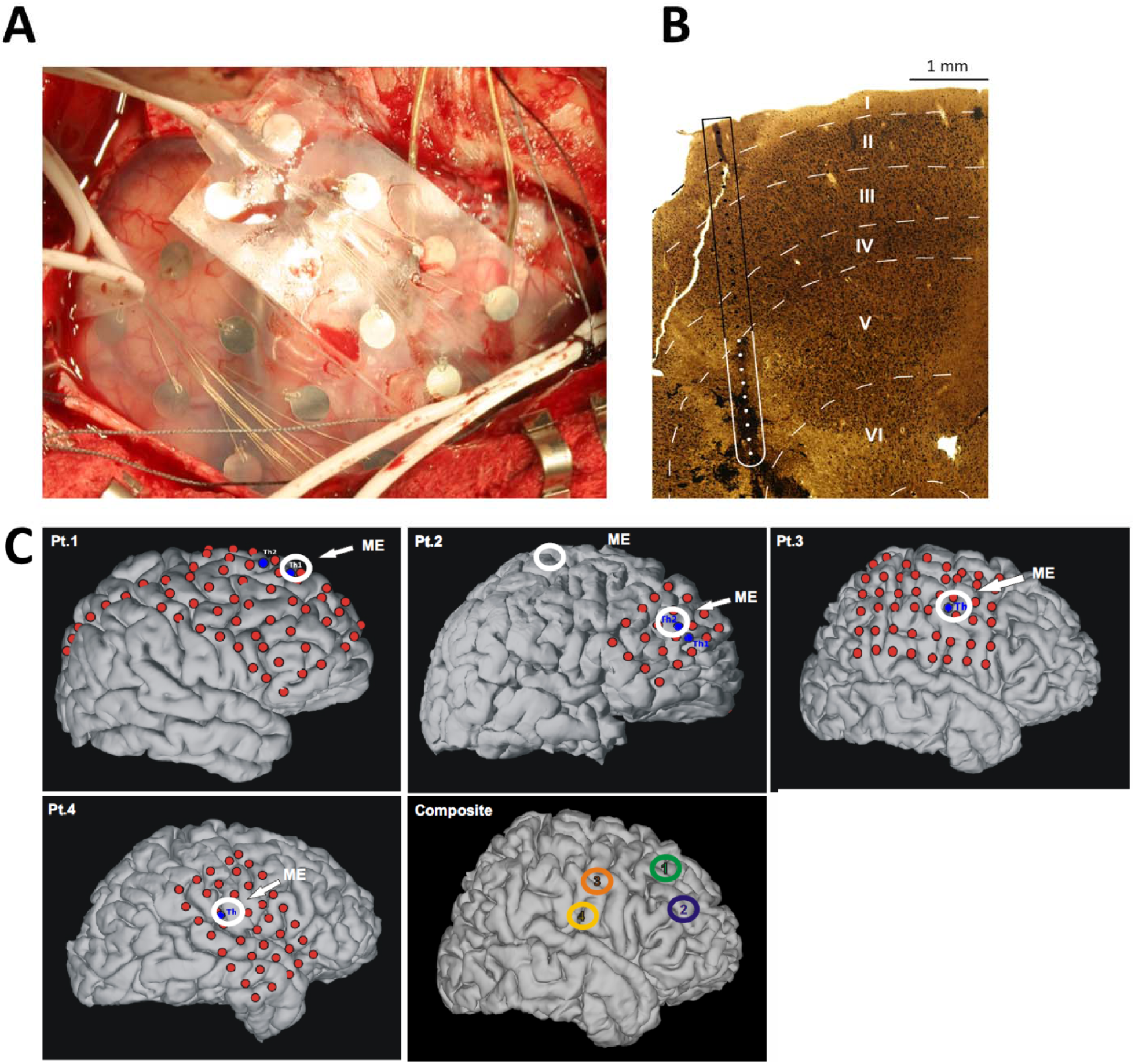
**Panel A:** ECoG and IME implantation procedure. **Panel B:** schematic representation of the localization of IME contacts with postoperative histology. The histological reconstruction of the tissue surrounding the IME in Patient 1 is shown. **Panel C:** postoperative MR reconstructions of the location of ECoG and IME electrodes from all subjects.

LFPg was equivalent to the data originally recorded from the IME, corresponding to the voltage difference between neighboring IME channels at a distance of 150 μm (μV/150 jam, henceforth referred to as μV). Current source density (CSD) was calculated as the negative of the second spatial derivative of the field potentials (Keller et al., 2009; Csercsa et al., 2010). However, since LFPg already constituted the first spatial derivative, CSD was calculated by simply calculating the negative of the first spatial derivative of the LFPg (Cash et al., 2009; Csercsa et al., 2010). Technical issues related to amplifier settings and file conversion resulted in raw IME signal voltage differences across subjects. Therefore, we generally z-transformed signal amplitude within subject for statistical analyses and indicate when it was otherwise. Unstandardized signal voltage, when reported, is not directly comparable across subjects.

Hypnograms were visually scored for ECoG data on a 20 second basis using standard criteria (Iber et al., 2007). Since the standard scoring criteria are generally applicable to scalp EEG channels with a full polysomnography setup (including EOG and EMG), we restricted our hypnograms to the identification of NREM sleep (regardless of stage) and the separation of it from other sleep states and wakefulness, based on the presence of slow waves and spindles (Ujma et al., 2015b). Artifacts were excluded from ECoG data on a 4 second basis using visual inspection. Only artifact-free data from NREM sleep was considered for further analysis.

ECoG and IME data were recorded separately, and unified again using a synchronization signal generated by E-prime 2.0 (Psychology Software Tools, Pittsburgh, PA), applied to the deepest IME contact (which was henceforth excluded from further analyses) and a randomly selected ECoG channel. The synchronization signal consisted of a unique sequence of marks which encoded the date and time at specified time intervals (10, 30 or 60 seconds). The unique sequences of synchronization marks were identified post-hoc in the simultaneously recorded ECoG and IME data, allowing for the synchronization of the two signals with data point precision.

### ECoG data analysis

ECoG data was analyzed using the Individual Adjustment Method (IAM) (Bódizs et al., 2009; Ujma et al., 2015a) in order to identify sleep spindles. The IAM essentially defines sleep spindles as waveforms with a specific frequency and sufficient amplitude that contributes to the spectral peaks of the NREM sleep EEG. In the first step, the high-resolution (0.0625 Hz) amplitude spectrum (AS) of the visually scored, artifact-free NREM sleep EEG was calculated for each ECoG channel free of interictal discharges (IIDs, “spikes”) based on 4 sec segments (Hanning-tapered, zero padded to 16 sec in order to ensure sufficient resolution in the frequency domain). The second-order derivatives of the averaged AS were calculated in order to identify spectral peaks. The resulting second-order derivatives were averaged across all IID-free ECoG channels. (This was a deviation from the original IAM methodology which averages second-order derivatives across frontal and centro-parietal scalp channels separately. This change was necessary because ECoG arrays did not always contain both frontal and centro-parietal channels.) A spectral peak was defined as the interval within which second-order AS derivatives were below zero. In line with the assumptions of the IAM about the spectral characteristics of sleep spindles, two clear spectral peaks were visible for each subject, one for slow spindles and another for fast spindles. The areas between these spectral peak boundaries across the frequency domain were defined as the slow and fast sleep spindle frequency range of the subject, respectively. In the second step, ECoG data was filtered to the subject’s individual slow or fast sleep spindle frequency range, and sleep spindles were identified as events in which the amplitude exceeded an electrode-specific threshold defined as the average of the AS values at the spectral peak boundaries, multiplied by the number of high-resolution frequency bins within the frequency range. If this dynamically defined threshold was not exceeded for at least 0.5 second, no sleep spindle was detected.

Sleep spindles were detected in this manner for IID-free ECoG channels and IME channels with no permanent artifacts. An ECoG spindle was considered as “local” if it was detected in less than or equal to 33% of the IID-free ECoG channels, and “global” if it occurred in more than 33% of IID-free ECoG channels. This cutoff point was chosen to ensure a similar number of local and global spindles (Table 1), but analyses confirmed the similarity of the laminar profile of all spindles (see Results, especially Figure 8), rendering the precise definition of the local-global cutoff point irrelevant.

**Table 1.** Basic clinical parameters and ECoG spindle counts by subject. The name of the drugs used in anti-epileptic medication regimens are abbreviated; CBZ: carbamazepine, CLON: clonazepam, LEV: levetiracetam, LTG: lamotrigine, OXC: oxcarbazepine, PHT: phentoin.

|  | Patient 1 | Patient 2 | Patient 3 | Patient 4 |
| --- | --- | --- | --- | --- |
| Age | 34 | 15 | 12 | 19 |
| Sex | Female | Male | Male | Male |
| MRI finding | Normal | Right frontal dysgenesis | Right parietal dysgenesis | Right fronto-centro-opercular dysgenesis |
| Seizure type | Focal, tonic postural | Focal, tonic postural | Focal, sensory-motor hemiconvulsive | Focal hypermotor |
| Pharmacotherapy | OXC, LEV, CLON | PHT, LTG | CBZ, OXC | CBZ, LTG, PHT |
| IME location | Right superior frontal gyrus | Right frontal | Right gyrus precentralis | Right gyrus precentralis |
| Slow spindle frequency range (Hz) | 11.45-12.61 | 10.43-11.54 | 12.26-13.5 | 11.28-12.22 |
| Fast spindle frequency range (Hz) | 13.3-14.67 | 12.49-13.38 | 13.64-15.28 | 13.49-14.44 |
| N<sub>slow, global</sub> | 73 | 335 | 34 | 404 |
| N<sub>slow, local</sub> | 19 | 256 | 97 | 167 |
| N<sub>fast, global</sub> | 76 | 460 | 28 | 324 |
| N<sub>fast, local</sub> | 19 | 356 | 84 | 241 |

The exact timing of ECoG sleep spindles were defined on the channel spatially closest to the IME for the analysis of laminar dynamics. ECoG data from this channel was filtered to the individual slow and fast spindle frequency, respectively, as it was determined in the first step of the IAM (two-way FIR filtering). IME data was triggered to the maximum positive deflection of the spindle cycle closest to the point where the envelope (defined as the modulus of the Hilbert transform of the filtered signal) of the filtered signal of this spatially closest ECoG electrode was maximal (spindle peak).

IME LFPg data segments from −500 msec to 500 msec relative to the identified ECoG spindle peaks were selected for further analysis. All selected data segments were visually inspected again in order to identify artifacts which were not present in the ECoG data. Artefactual data segments were excluded. All data segments were filtered to a generic sleep spindle range (10-16 Hz, two-way FIR filtering implemented in Matlab EEGLab) and smoothed across layers using a Hamming window (Ulbert et al., 2001; Csercsa et al., 2010). CSD was calculated by data segment after selection, filtering and smoothing.

In order to quantify laminar differences in field potential and current strength, respectively, the root mean square (RMS) of the LFPg and CSD signal of each channel was calculated for each 1-second IME data segment. Channel-wise RMSs were averaged across channels in the same cortical layer, yielding an average of LFPg and CSD signals for each spindle and for each cortical layer (LFPg/CSD magnitude).

### Multiple unit and single-unit activity

Multiple unit activity (MUA) was estimated by calculating the filtered (300-3000 Hz, zero phase shift, 48 dB/octave) and rectified LFPg sampled at 20 kHz, and passing it through a low-pass filter (20 Hz, 12 dB/octave) (Csercsa et al., 2010). MUA was triggered to ECoG spindle detections in a similar manner as IME data from the EEG range and averaged across detections by spindle type (slow/fast, local/global). MUA was only computed from the two patients (Patient 1 and 2) with adequate high-sampling rate signal quality.

Single-unit activity (SUA) was performed on high sampling rated IME recordings (20kHz) in case of two patients (Patient 1 and 2), where individual cell activities could be separated reliably from MUA/field activity and background noise. SUA was performed on epoched raw IME data sampled at 20 kHz, initially extracting two second epochs (±1000 msec around all ECoG spindle peaks). We epoched and analyzed data from slow, fast, local and global spindles separately. After DC offset removal, we detected SUA based with an amplitude threshold adjusted manually according to the magnitude of background noise of each channels. Multiple individual neurons were identified as the generators of SUA on each channel based on clustering by action potential morphology and amplitude in a 0.4 msec timeframe. We applied a principal component analysis based on waveform characteristics to refine clustering and reduce false detections. All SUA detections were performed in Spike2 (version 7 software (CED Limited, UK). SUA occurrence relative to LFPg phases was calculated based on the instantaneous phase angle of LFPg signal’s Hilbert transform. All SUA detections within ±500 msec from ECoG spindles of the same spindle type were pooled from all cells on the same channel and from all IME channels in the same layer, while the rest more temporally removed from ECoG spindle peaks were discarded.

### Data analysis software

EEG analysis was performed using Fercio© (Ferenc Gombos, Budapest, Hungary) for the IAM analysis, MATLAB 2018b (The MathWorks, Inc., Natick, MA) EEGLab (Swartz Center for Computational Neuroscience, San Diego, CA) for filtering, envelope calculation and data synchronization, and Neuroscan (Compumedics USA, Charlotte, NC) for the selection and processing of LFPg and CSD segments. Statistical analysis was performed in STATISTICA 10.0 (StatSoft, Tulsa, OK) and custom scripts using MATLAB 2018b. The statistical analyses of SUA were performed using Oriana (v2.02c, Kovach Computing Services).

## RESULTS

### Sleep spindles on the IME

In the first step, we obtained basic information about the occurrence of spindle-like oscillations on IME recordings. Transient, waxing and waning oscillations with morphological characteristics resembling EEG and ECoG sleep spindles were clearly visible in the IME recordings of 2 out of our 4 patients (Patient 1 and 2). In these patients, characteristic spectral peaks in the sigma range (8-16 Hz) also emerged once power spectral density was calculated from the IME signal, and these spectral peaks corresponded to those independently calculated from the ECoG signal. In two further patients, neither visible spindle-like IME oscillations nor characteristic spectral peaks emerged. However, even in these patients, once IME recordings were triggered to the peak of the spindles detected on the nearest ECoG electrode, a clear spindle-frequency oscillation emerged in the averaged signal, indicating that a sub-threshold spindle-frequency signal was always concomitant to ECoG spindles **(Figure 2).**

**Figure 2.**
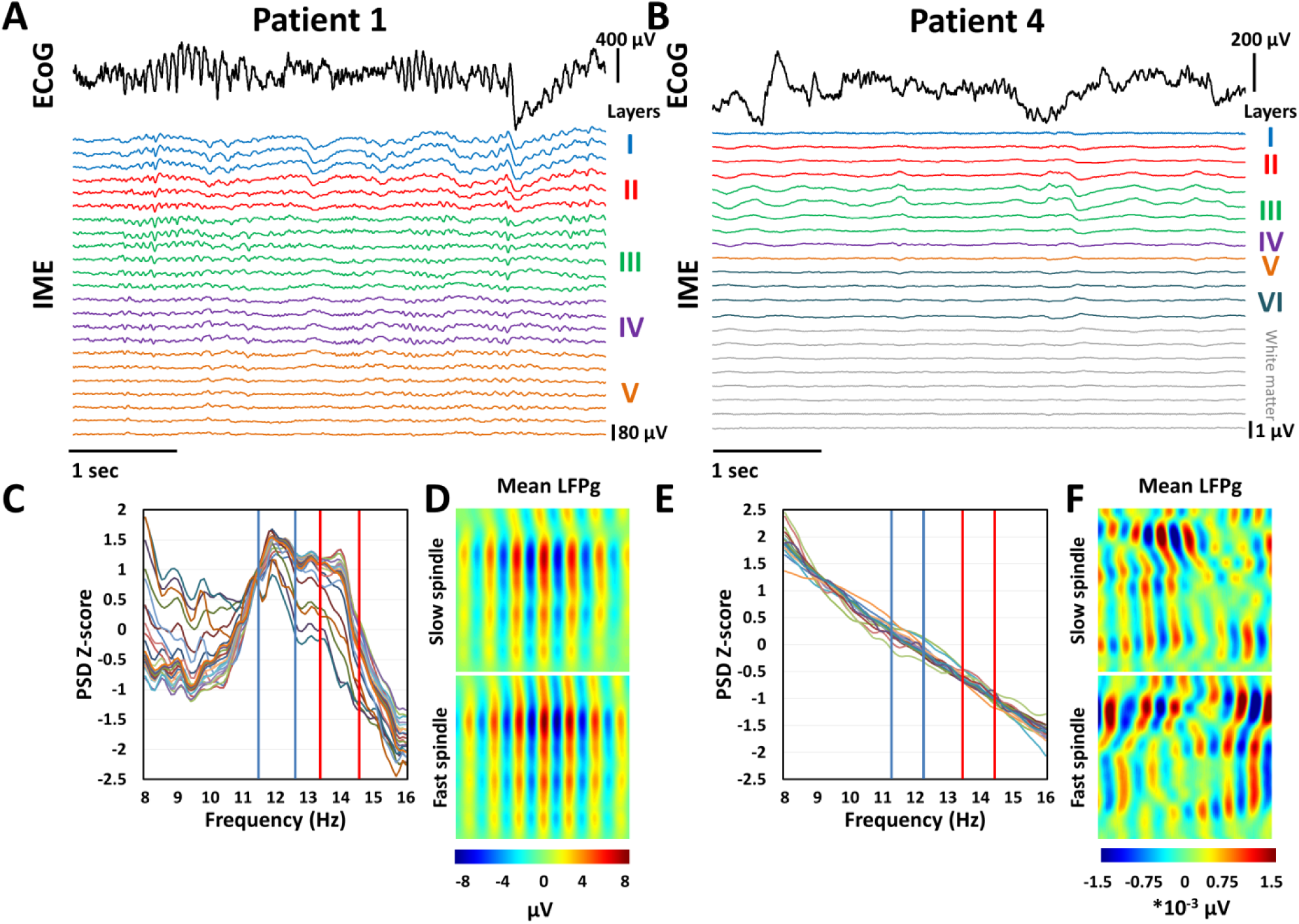
Sleep spindles are not always visible on IME LFPg recordings and do not always create characteristic spectral peaks, but they always emerge when ECoG spindle detection-triggered IME LFPg data is averaged. In Patient 1, both slow and fast spindles visible on ECoG recordings are also visible in IME derivations **(Panel A)**, and spectral analysis reveals that the spectral peaks identified from ECoG are also present in IME data **(Panel C**, showing power spectral density from every IME channel after z-transformation by channel. Blue and red vertical lines represent the slow and fast sleep spindle frequency ranges, respectively, as determined from ECoG data). Averaged IME LFPg triggered to ECoG spindle detections also reveals the presence of spindles on the IME during ECoG spindles **(Panel D).** In Patient 4, however, sleep spindles are not visible present in the raw IME data during ECoG spindles **(Panel B)**, do not leave characteristic spectral peaks **(Panel E)**, but appear when averaged IME LFPg is triggered to ECoG spindle detections **(Panel F).** Note that IME voltage values are not directly comparable across subjects due to technical issues resulting in large differences in mean voltage.

### Sleep spindle co-occurrence across ECoG and IME channels

Since in two patients the IME signal was characterized by spindle-like oscillations and sigma frequency peaks at the same frequency where they were observed in the ECoG signal, we proceeded to investigate the general pattern of occurrence of micro-domain IME spindles as well as their co-occurrence with macro-domain ECoG spindles. We performed a direct detection of sleep spindles in the IME recordings of the two patients whose recording contained clearly visible spindles and sigma spectral peaks, and compared these to the sleep spindles detected from the ECoG signal of the same patients recorded at the same time.

No ECoG electrode or cortical layer was without spindle detections. In ECoG detections, spindle density exhibited a slight variation across cortical locations, in line with previous findings (Andrillon et al., 2011; Peter-Derex et al., 2012; Piantoni et al., 2017) **(Figure 3, Panel A).** In direct IME detections, sleep spindle density was relatively uniform across layers **(Figure 3, Panel B)**, however, amplitude was higher in the superficial layers **(Figure 3, panel C).** In both ECoG and IME detections, the majority of sleep spindles occurred on a low number of ECoG channels and in only one cortical layer, respectively, and more global spindles were progressively rarer **(Figure 3, Panel D-E).** (Note that the proportion of global and local spindles on the ECoG, discussed in later in the Results, is not directly related to this finding since for the laminar analysis we only considered local spindles which occurred on the specific ECoG electrode closest to the IME). Amplitude was positively associated with the number of channels/layers a spindle was detected on for both ECoG and IME **(Figure 3, Panel F-G).** Density and amplitude measures are reported here as the mean density and amplitude, respectively, of all IME channels within a cortical layer.

**Figure 3.**
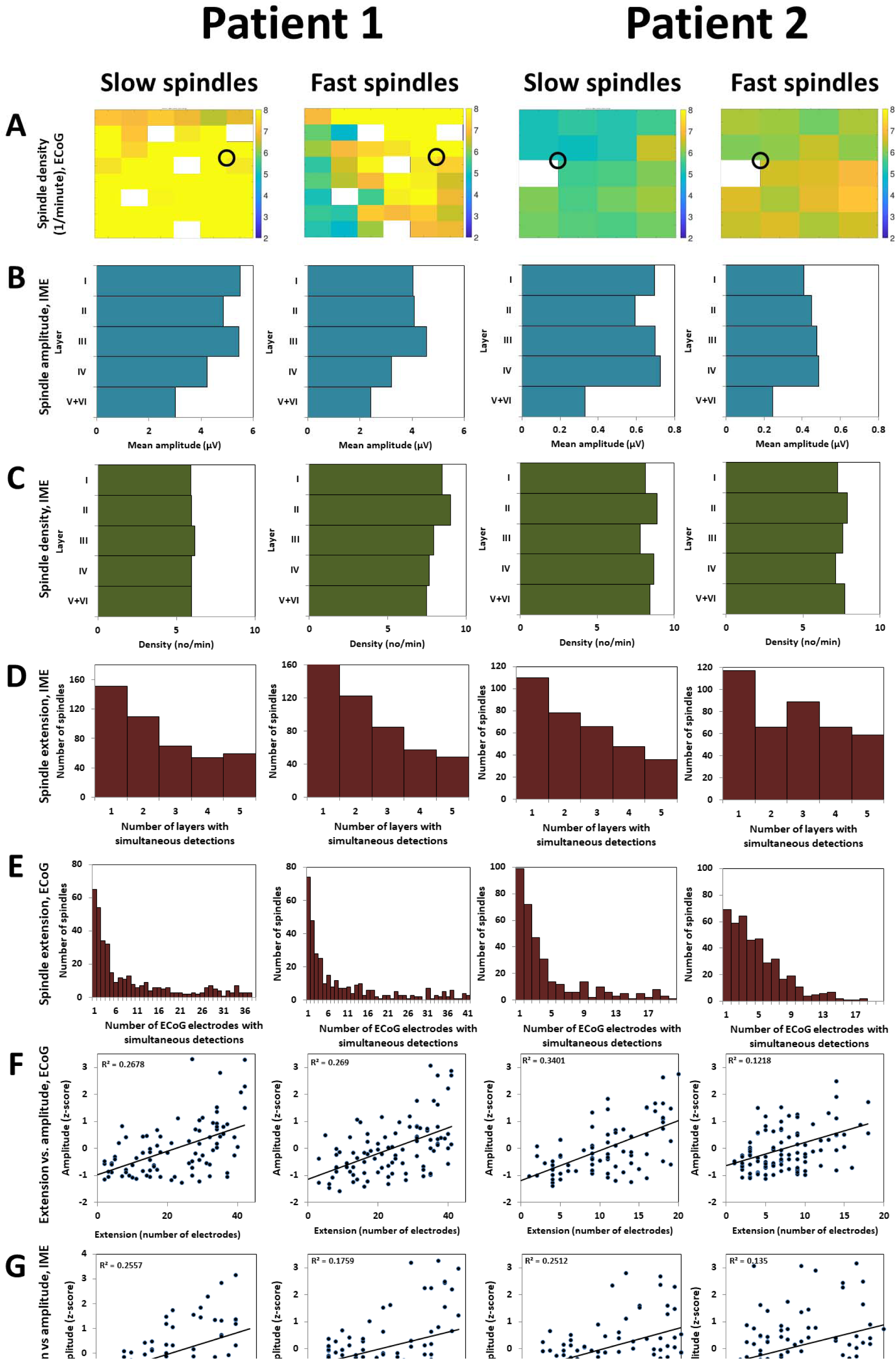
**Panel A:** sleep spindle density on the ECoG. The color matrix is a topographically accurate representation of the implanted electrode grid of each patient. Electrodes with unusable or pathological data are left pure white. The location of the IME is marked with a circle. **Panel B:** sleep spindle amplitude on the IME. For each layer, sleep spindle amplitude is given as the mean value of all channels in the layer. **Panel C:** sleep spindle density on the IME. For each layer, sleep spindle density is given as the mean value of all channels in the layer. **Panel D:** histogram of the spatial extent of sleep spindles on the ECoG, given as the number of spindles (axis y) co-occurring on a given number of ECoG channels (axis x). The first 20 bins (this number is limited by the lower number of ECoG electrodes in Patient 2) of the four histograms are not significantly different (2-sample Kolmogorov test, p_min_=0.27 from all possible comparisons). **Panel** E: histogram of the spatial extent of sleep spindles on the IME, given as the number of spindles (axis y) simultaneously detected in a given number of cortical layers (axis x). Only spindles occurring on at least **1/3** of the channels in the same cortical layers were considered. The four histograms are not significantly different (2-sample Kolmogorov test, p_min_=0.69 from all possible comparisons). **Panel F:** the association between ECoG sleep spindle amplitude (axis y) and extension defined as the number of ECoG electrodes with simultaneous detections (axis x). Data is shown for the ECoG channel closest to the IME. Correlation coefficients are not significantly different (Fisher’s r-to-z method, p_min_=0.23 from all possible comparisons). **Panel G:** the association between IME sleep spindle amplitude (axis y) and extension defined as the number of IME electrodes with simultaneous detections (axis x). Data is shown from a representative channel with high spindle density. In Patient 2, slow spindle amplitude is significantly more strongly correlated with extension than fast spindle amplitude **(p=0.044)**, all other correlations are not significantly different (Fisher’s r-to-z method, p_min_=0.13 from all other possible comparisons).

In the co-occurrence analyses, we considered two spindle events from two electrophysiological sources s1 and s2 to co-occur if a sleep spindle detected in s2 started before a concomitant sleep spindle in si terminated, s1 always indicating the channel where the earlier spindle was detected. For cortical layers, only spindles co-occurring on at least 1/3 of the channels within the cortical layer using the above criteria were considered in order to control for the higher prior probability of a sleep spindle occurring at a given time (and thus co-occurring with others) as a function of the number of channels within the same layer. As for the co-occurrence between IME and ECoG, a sleep spindle was considered to co-occur if it occurred simultaneously using the above criteria in any cortical layer and an ECoG electrode. A large proportion of sleep spindles co-occurred between cortical layers (typically 50-80%) and IME and ECoG (typically >90% of ECoG spindles co-occurring on the IME). Between-layer co-occurrence was most common between neighboring layers and IME-ECoG co-occurrence was less common as a function of distance between ECoG channel and the IME. IME-exclusive spindles were relatively common, but most ECoG spindles were accompanied by spindles on the IME (Figure 4).

**Figure 4.**
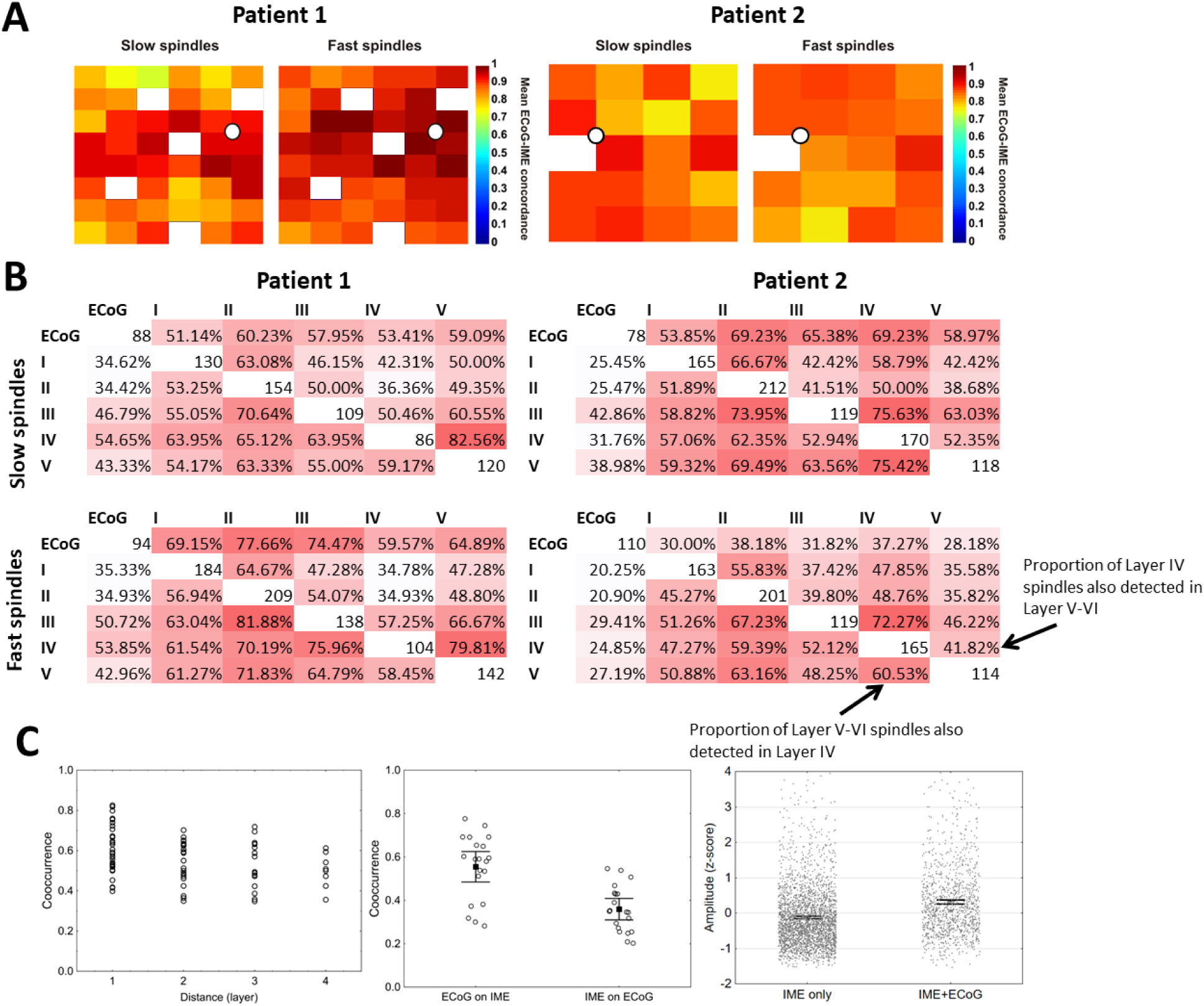
**Panel A:** the proportion of ECoG spindles with a concomitant IME spindle in at least one cortical layer. The color map is a topographically accurate representation of the implanted subdural electrode grid with a white dot indicating the location of the IME. Electrodes with unusable or pathological data are colored pure white. **Panel B:** the number and proportion of ECoG spindles co-occurring between specific cortical layers mapped by the IME and the nearest ECoG channel in Patient 1 and Patient 2. Only sleep spindles simultaneously detected on at least 1/3 of the IME channels within the same cortical layer were considered. The diagonals indicate the number of spindles occurring in each cortical layer or on the nearest ECoG channel. The upper half of the matrix shows the proportion of spindles in the source indicated on axis y co-occurring with spindles in the source indicated on axis x, while the lower half shows the proportion of spindles in the source indicated on axis x co-occurring with spindles in the source indicated on axis y. An example of this interpretation is provided on the figure. **Panel C, left:** the negative relationship between source distance (in layers) and the probabilities of co-occurrence depicted in Panel B (Pearson’s r=-0.31, p=0.005). All spindle types from both patients were pooled for this analysis, within-patient and within-type correlations range between −0.28 and −0.37. **Panel C, middle:** The probability of ECoG spindles co-occurring on the IME within a cortical layer (ECoG on IME) is higher than the probability of IME spindles within a cortical layer co-occurring on the ECoG (IME on ECoG) (One-way ANOVA F=22.65, p<0.001). The box and whiskers show means and 95% confidence intervals, individual data points (co-occurrence probabilities between ECoG and each layer from Panel B) are shown. Data from all layers, patients and spindle types were pooled for this analysis. **Panel C, right:** individual IME spindles that co-occur on ECoG have higher amplitude than those that do not (F=166.6, p<0.001). Data from all layers, patients and spindle types were pooled for this analysis, amplitudes are z-transformed by patient, layer and spindle type. The box and whiskers show means and 95% confidence intervals, individual data points except 30 spindles with standardized amplitude>4 are shown. This pattern is present in all layers (F=13.9-48.2, p<0.001 in all cases).

These results were not substantially different if we only considered spindles which co-occurred in *1/2* or 2/3 of the IME channels within the same layer, apart from a reduction in the total number of co-occurring spindles.

We hypothesized that the simultaneous appearance of sleep spindles on IME and ECoG recordings is more likely as a function of increasing amplitude: that is, sufficiently large IME spindles appear on the ECoG (or on more ECoG electrodes) while others do not. This hypothesis was supported by the fact that the correlation between IME spindle amplitude and the spatial extension of concomitant ECoG spindles is universally positive and statistically significant 13 out of 20 cases instead of the single case expected under a null hypothesis (Table 2). Note that a perfect correlation is not expected due to the variable degree of overlap across spindles between the actual cortical area of spindle occurrence and the cortical area where ECoG and IME are present.

**Table 2.** Pearson correlation coefficients between the amplitude of IME sleep spindles and the spatial extension (expressed in the number of ECoG electrodes the spindle was simultaneously detected on) of concomitant ECoG spindles. Results are presented by the cortical layer of IME spindle detection, patient and spindle type.

|  |  | Layer |  |  |  |  |
| --- | --- | --- | --- | --- | --- | --- |
|  |  | I | II | III | IV | V+VI |
| Patient 1 | Slow | 0.163 | 0.098 | 0.059 | 0.205** | 0.333** |
|  | **Fast** | 0.066 | 0.242** | 0.162* | 0.371** | 0.268** |
| **Patient 2** | **Slow** | 0.388** | 0.529** | 0.475** | 0.494** | 0.308** |
|  | **Fast** | 0.042 | 0.230** | 0.137* | 0.078 | 0.077 |

\* p<0.05
\*\*p<0.01

### Spindle type effects on IME spindles

So far we have established that micro-domain spindles detected on the IME are characterized by a highly heterogeneous cortical profile, and various patterns of spindle occurrence can be observed, in line with previous findings (Hagler et al., 2018). We next investigated whether the occurrence of sleep spindles directly detected from the IME is affected by spindle type (slow/fast, local/global): that is, whether different spindle types are preferentially detected in one or another layer, as previously hypothesized (Timofeev and Chauvette, 2013; Piantoni et al., 2016).

In order to test whether the occurrence of slow and fast spindles is different at various cortical depths, we compared the density of slow and fast spindles across all layers. The density of slow and fast spindles was not significantly different in any layer, (X^2^-test with row-wise z-tests, all p>0.05, **Figure 3, Panel B).**

We also assessed whether the layer of occurrence for IME spindles is affected by ECoG spindle globality - for example, whether IME spindles occurring in more superficial layers co-occur with more global ECoG spindles. (Note that while slow and fast spindles could be identified from the IME alone, spindle globality could only be assessed based on co-occurring ECoG spindles because only the ECoG spanned a substantial area of the cortex, making it possible to assess how widespread a spindle was). We treated ECoG spindle globality as a continuous variable defined as the number of ECoG electrodes a spindle was detected on, and assessed whether IME spindles detected in any layer usually co-occur with more global ECoG spindles. This was not the case: co-occurring ECoG spindle globality was not significantly different between any two layers, with one exception: in Patient 2, sleep spindles detected in Layer V/VI co-occurred with significantly less global ECoG spindles than those in other layers **(Figure 5).**

**Figure 5.**
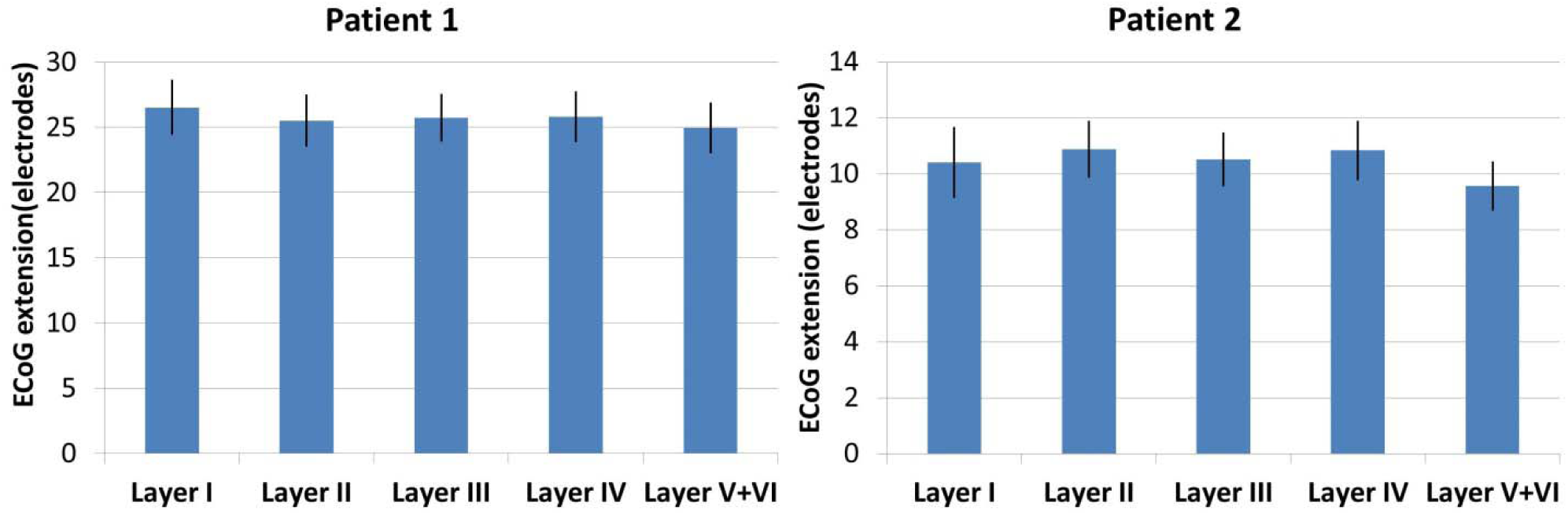
The extension (number of ECoG channels a spindle was detected on) of ECoG spindles co-occurring with IME spindles detected from each cortical layer in Patient 1 **(Panel** A) and Patient 2 **(Panel B).** Error bars denote 95% confidence intervals. Slow and fast spindles were pooled for this analysis.

Therefore, it can be stated that the laminar profile (or more precise, layer of occurrence) of micro-domain IME spindles is heterogeneous, but similar for slow, fast, local and global spindles.

### The laminar profile of ECoG sleep spindles

Since we have established that at least sub-threshold spindle oscillations were present in the IME recordings of all patients, we next investigated whether the laminar profile - that is, the characteristic pattern of LFPg and CSD amplitude across layers - of macro-domain spindles detected on the ECoG is affected by type (slow/fast) or globality (global/local). As opposed to the micro-domain spindle detectable by the IME recordings, ECoG spindles can be considered macro-domain spindles due to the larger receptive fields of ECoG electrodes. We detected sleep spindles in all physiological ECoG channels, and triggered the signal of IME channels to the peaks of ECoG spindles detected from the ECoG electrode closest to the IME. We distinguished between slow and fast spindles (based on the frequency ranges determined by the IAM method, see Methods) and local and global spindles (based on whether or not the spindles were present on more or less than 33% of all physiological ECoG channels). The effects of sleep spindle characteristics on the laminar profile were estimated using a mixed-effects ANOVA model using Subject, Spindle type (slow/fast) and Globality (local/global) as between-subject factors, Layer as a within-subject factor, and Magnitude (defined as the root-mean-square of the IME signal in the 500 msec preceding and following the ECoG spindle peak, averaged across IME channels within the same layer) as the dependent variable. This model was run separately for the two transformations of the IME signal: local field potential gradients (LFPg magnitude) and population transmembrane currents (CSD magnitude). Due to the great statistical power resulting from the large number of spindles (N=2973) we also report the effect size of test statistics, expressed as the partial eta square (η^2^) which is more indicative than pure statistical significance. η^2^ is a measure of the proportion of variance explained by a main effect or interaction, controlling for the effects of other predictors thus not biasing the estimate downwards in case of many predictors (Richardson, 2011). We report the results of this analysis in **Supplementary Tables 1-2.** In short, large between-subject differences in the raw magnitude and laminar profile distribution of sleep spindles were observed, while Layer, Spindle type and Globality had an intermediate effect size and more superficial layers, slow spindles and global spindles were associated with larger LFPg and CSD magnitude. The interaction effects of Spindle type*Layer and Globality*Layer, indicating a different distribution of magnitude across cortical layers in global/local and slow/fast spindles, were sometimes significant but very modest in effect size (partial η^2^_max_=0.01), indicating that at least within the same patient slow spindles, fast spindles, local spindles and global spindles were characterized by a similar laminar profile. These effects are illustrated in **Figure 6.**

**Figure 6.**
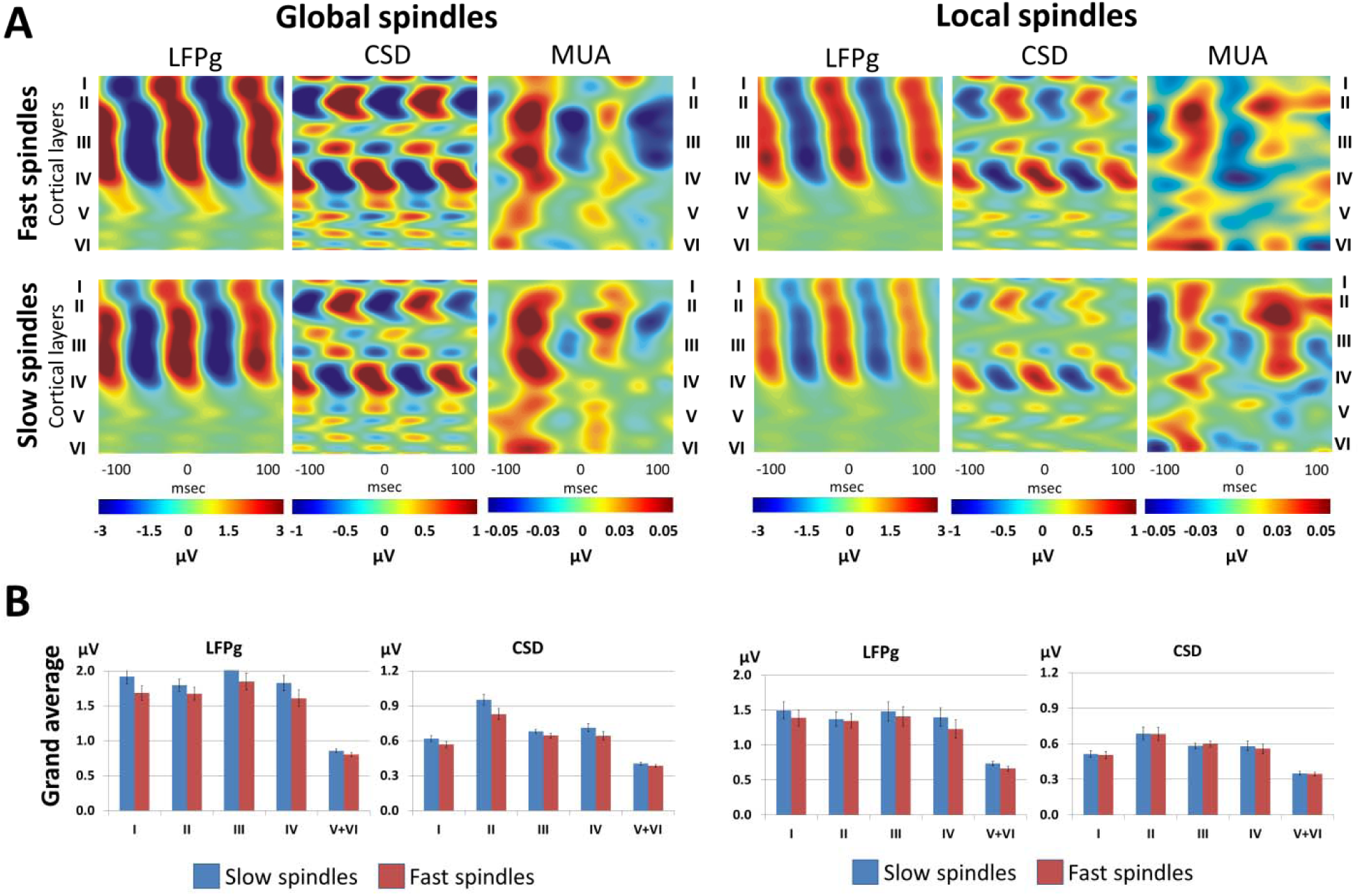
**Panel A:** The laminar profile of slow, fast, local and global spindles, including LFPg, CSD and MUA, in representative Patient 2. Color maps illustrate the average amplitude fluctuations ±100 msec before and after ECoG spindle peaks. Layer centroids are marked with Roman numerals. Note the similarity of the profile across all spindle types. **Panel B:** Grand average LFPg and CSD magnitude across layers for slow and fast spindles from all patients. μV values refer to μV/150 μm in case of LFPg and μV/150 μm^2^ in case of CSD.

These analyses were repeated using z-transformed LFPg and CSD magnitudes across layers in order to eliminate the potential bias of between-subject differences in mean voltage. This approach is unable to yield between-subject main effects, as the mean magnitude across layers for each spindle is set to 0 with a standard deviation of 1, but it is sensitive to the interactions between the within-subject factor Layer and between-subject factors, and thus sensitive to the modification of the laminar profile by between-subject factors such as spindle type or spindle globality. The results were very similar to the non-normalized model: between-subject heterogeneity was substantial, but the laminar profile of all spindle types was very similar within the same subject (interaction partial η^2^_max_=0.007). Detailed results are provided in **Supplementary tables 1-2.** The between-subject heterogeneity but within-subject homogeneity - regardless of spindle type - of the laminar profile of spindles is illustrated on Figure 7.

**Figure 7.**
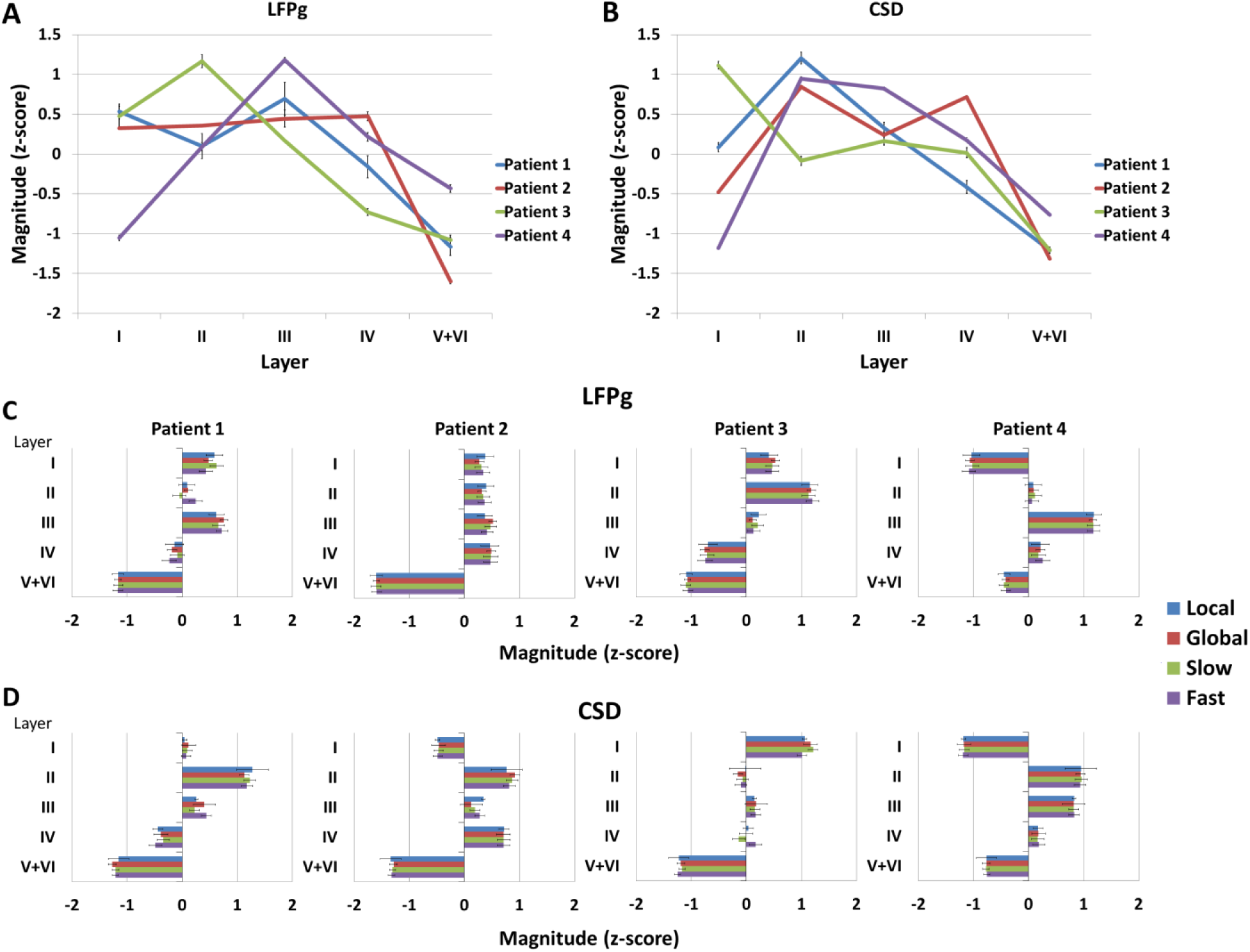
Normalized LFPg **(Panel** A and **C)** and CSD **(Panel B** and **D)** laminar profiles of sleep spindles. **Panel** A and **B** illustrate the mean LFPg and CSD magnitude of each subject in each layer, respectively. **Panel C** shows LFPg magnitude in each layer for each subject by spindle type (local, global, slow and fast), while **Panel D** shows the same statistics for CSD magnitudes. Error bars represent 95% confidence intervals on all plots. Note the similarity of the individual laminar profile across spindle types.

In order to confirm LFPg and CSD magnitude during ECoG spindles really follows a similar pattern across layers regardless of spindle type (slow/fast or local/global), we z-transformed both LFPg and CSD amplitude within each layer of each patient across all detections, and performed a principal component analysis on the magnitude values in each layer. In each patient, only one principal component with an eigenvalue>1 emerged, explaining 62-82% of the variance in LFPg magnitude and 60-76% of the variance of CSD magnitude across layers. The high multicollinearity of LFPg and CSD amplitude across cortical layers is illustrated in **Figure 8.** A more illustrative, rotating version of this figure is provided in **Supplementary video 1** and **Supplementary video 2**. The similar signal amplitude across all layers observed in all spindles also suggests that the general similarity of local and global spindles we report are not specific to the 33% cutoff criterion.

**Figure 8.**
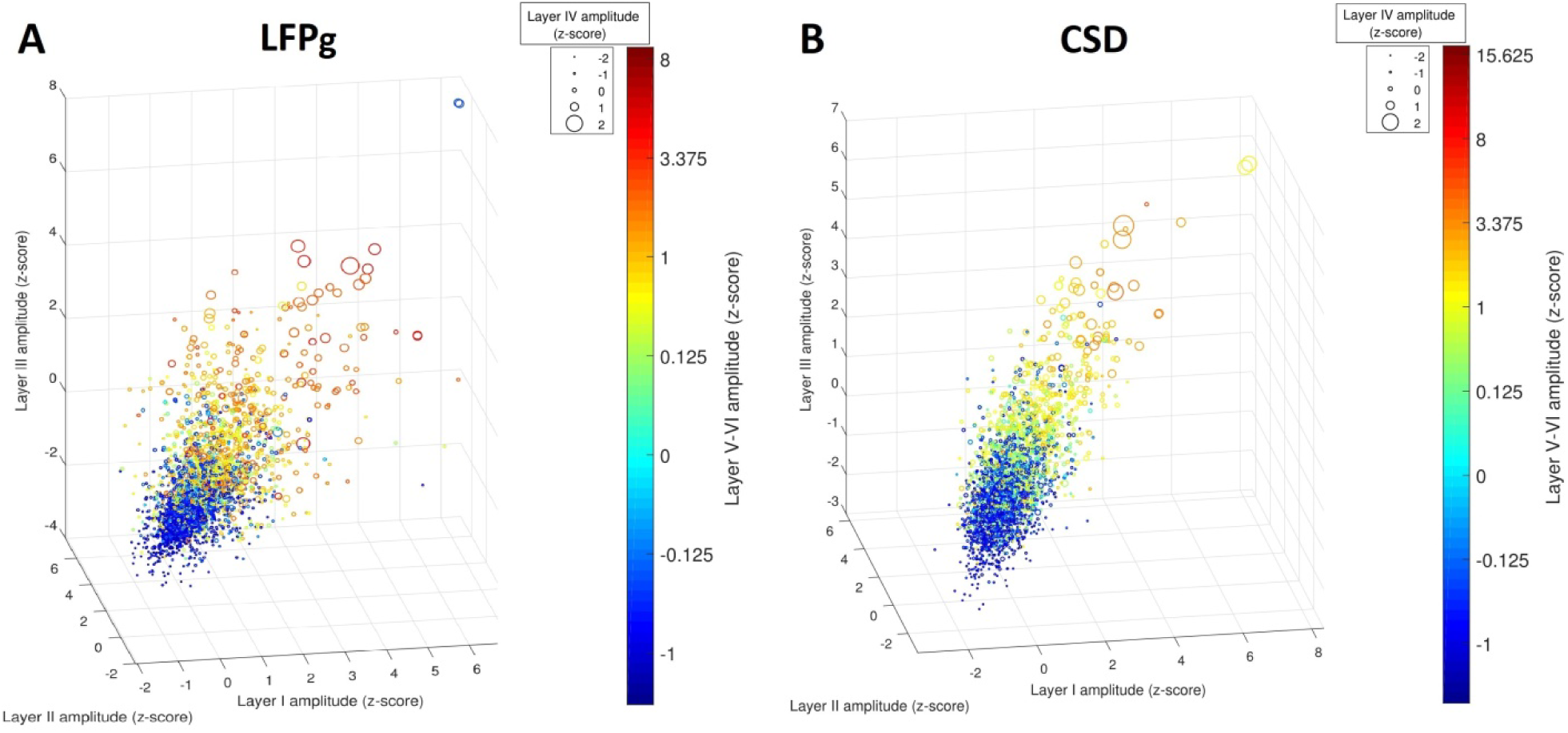
Scatterplot of LFPg **(Panel** A) and CSD **(Panel B)** magnitudes in each cortical layer during ECoG spindles. The three spatial dimensions are used to illustrate magnitudes in layer I-III, while marker sizes and colors illustrate magnitudes in layer IV and V/VI, respectively. Marker sizes and marker colors are transformed to nonlinear scales to optimize visibility.

In sum, all ECoG spindle types (slow/fast, local/global) were characterized by a similar pattern of signal amplitude in cortical layers, making it unlikely that neuronal networks with substantially different cortical innervation patterns contribute to different spindle types.

### Single- and multiple-unit activity

High-quality single-unit (SUA) and multi-unit (MUA) activity was obtainable from two patients. Based on a statistically significant deviation from circular uniformity, in Patient 1 SUA was significantly coupled to LFPg during all spindle types and in all layers except slow local, fast local and slow global spindles in Layer II and during fast local spindles in Layer V. In Patient 2, both slow and fast global spindles were coupled to LFPg in Layer II and III. Slow local spindles in Layer II and fast global spindles in Layer VI were also significantly coupled. In case of significant coupling, SUA preferentially occurred during positive-negative LFPg transitions in all patients **(Figure 9). Supplementary figures 1 and 2** illustrate the phase relationship between SUA and LFPg by patient, layer and spindle type using rose histograms.

**Figure 9.**
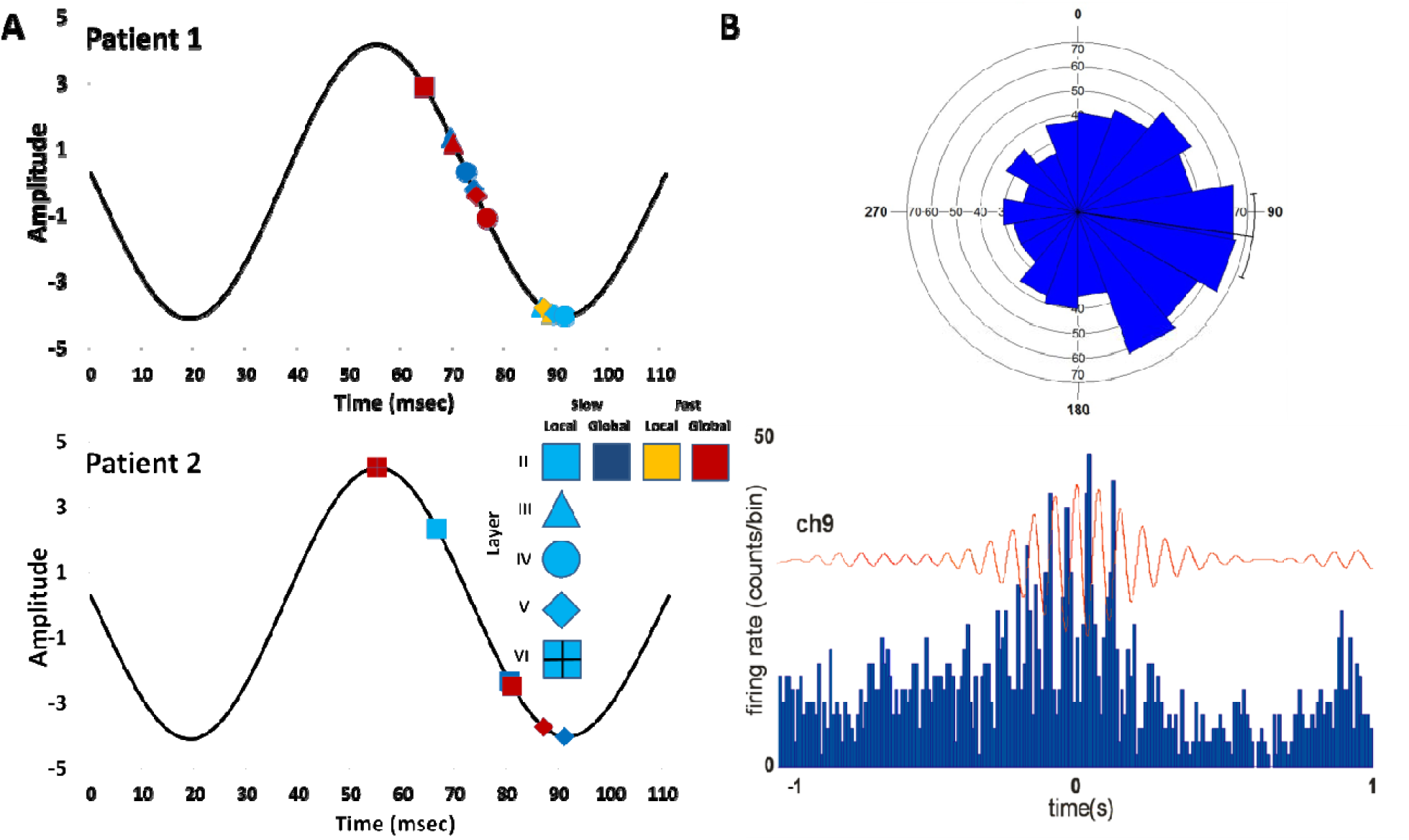
**Panel A:** the preferred phases of SUA during various spindle types, superimposed to a stereotypical spindle-frequency oscillation. Only those spindle types are shown for which LFPg-SUA coupling (based on an at least nominally significant Rayleigh’s Z-test for circular uniformity) was significant. Spindle type is coded by marker colors and layer of occurrence is coded by marker shapes. **Panel B:** an illustration of the occurrence of SUA during positive-negative LFPg phase transitions using data from fast global spindles and a layer III channel in Patient 1. The rose plot illustrates the preferred firing phase slightly after 90° (see also the first data marker on panel A). The line-histogram shows the frequency of occurrence of SUA in temporal bins superimposed to the mean spindle LFPg on the same channel.

A circular factorial ANOVA using the Harrison-Kanji test (Harrison et al., 1986) revealed a significant omnibus effect of both layer and spindle type on the preferred phase of SUA-LFPg coupling in both patients (all p<0.001). As post-hoc tests, we compared the preferred phases of SUA-LFPg coupling across layers and across spindle types using Mardia’s test of the difference of circular means (Mardia, 1969). In Patient 1, slow and fast spindles within the same layer typically had similar preferred phases, which were however different for the same spindle types in different layers, and for local and global spindles even in the same layer. Similar trends were seen in Patient 2, but they only reached significance when comparing superficial and layer V-VI global spindles. However, it must be noted that the statistical power of all comparisons is higher in Patient 1 due to more SUA events. Preferred SUA phases are listed in Table 3. Detailed statistics are provided in the Supplementary data.

**Table 3.** Preferred LFPg phases of SUA by spindle type. The preferred phases of distributions significantly different from circular uniformity are set in bold. Note that in case of the rest means and their differences may not be meaningful due to the circular uniformity of phases.

| Layer | Patient 1 |  |  |  | Patient 2 |  |  |  |
| --- | --- | --- | --- | --- | --- | --- | --- | --- |
|  | Slow |  | Fast |  | Slow |  | Fast |  |
|  | Local | Global | Local | Global | Local | Global | Local | Global |
| II | 147.213° | 18.727° | 343.005° | **50.511°** | **54.067°** | **153.495°** | 121.598° | **123.574°** |
| III | **159.739°** | **92.08°** | **149.316°** | **89.901°** | 149.03° | **169.306°** | 164.061° | **150.524°** |
| IV | **147.097°** | **70.955°** | **158.527°** | **71.723°** | 224.569° | 183.848° | 248.283° | 195.217° |
| V | **166.859°** | **83.996°** | 149.205° | **100.808°** | 199.599° | 241.988° | 237.037° | 319.986° |
| VI |  |  |  |  | 354.699° | 326.885° | 3.87° | **355.796°** |

MUA and LFPg from each channel was triggered to ECoG spindle events and averaged across events. The similarity of the resulting average signals was assessed by computing their Pearson’s correlation coefficient, and their phase difference was estimated by the angular mean of the difference of the phase angle of their Hilbert transforms. LFPg and MUA during spindle events were generally highly similar and exhibited an approximately antiphase (9O°<ΔΦ<27O°) relationship (Figure 10). However, the LFPg-MUA phase angle difference was not constant during spindle events, especially in Patient 2, leading to a downward bias of correlation coefficients and the poor representativeness of the average phase difference.

**Figure 10.**
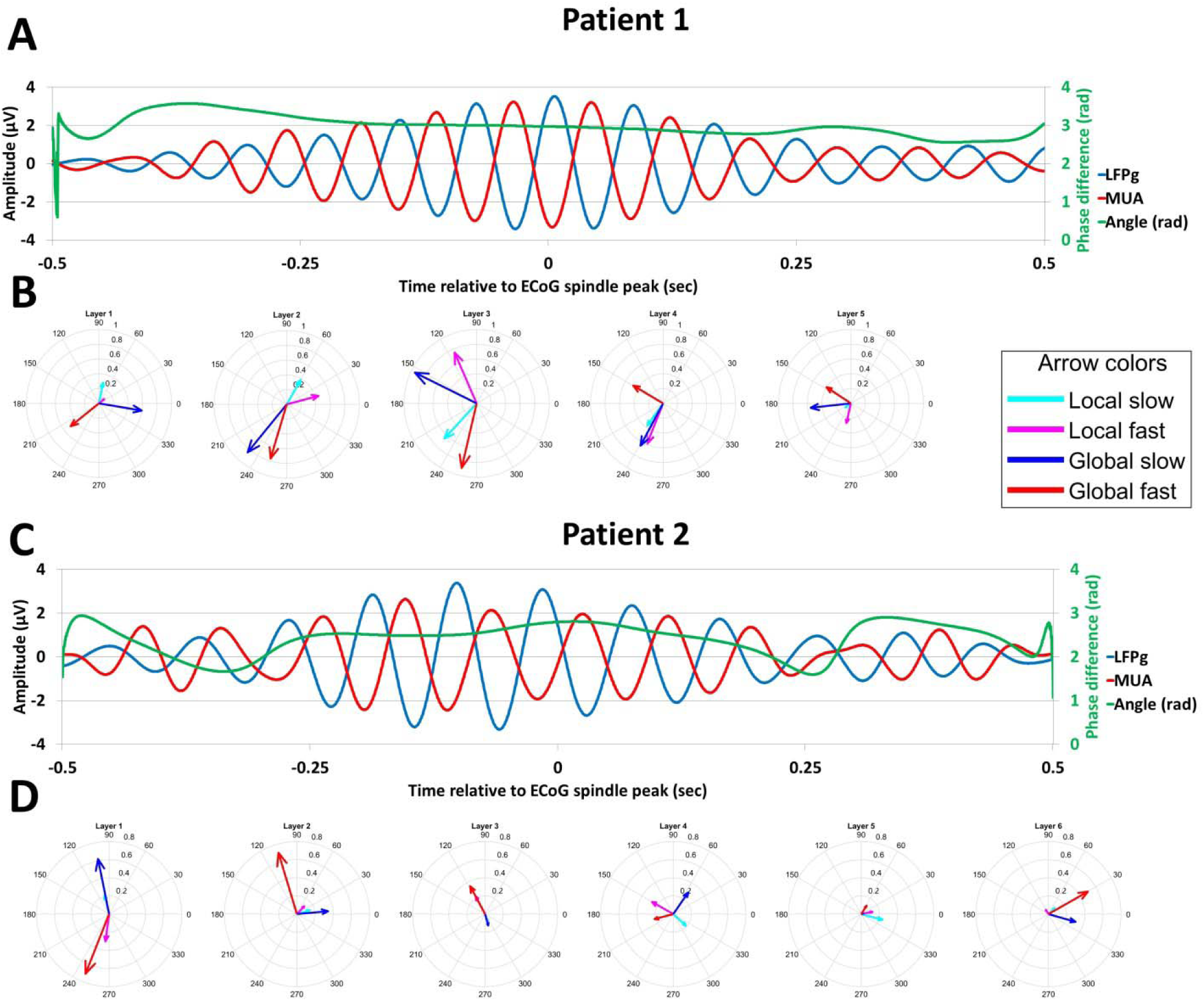
Representative LFPg and MUA averages, triggered to ECoG spindle events **(Panel A and C)** from channel 10 (Layer **III)** in Patient 1 and channel 6 (Layer **II)** in Patient 2. Note the fluctuations of the difference of instantaneous phase differences. **Panels B and D** illustrate the similarity of the LFPg and MUA signals as well as their phase difference by layer and spindle type in Patient 1 and 2, respectively. On the compass plots the length of the arrows corresponds to the modulus of the LFPg-MUA correlation coefficient, while the orientation of the arrows corresponds to the circular mean of the difference of the instantaneous phase of the two signals.

In sum, SUA and MUA was generally negatively correlated to LFP, but this relationship was not fully homogeneous across patients, layers and spindle types.

## DISCUSSION

In line with recent findings (Hagler et al., 2018), we found evidence for the presence of sleep spindles not only across the cortical mantle using ECoG (macro-domain spindles), but also on IME electrodes penetrating the cortex and recording electrophysiological activity within cortical layers (micro-domain spindles). Not all patients showed visible sleep spindles typical for ECoG and scalp EEG derivations on their IME recordings, but the ECoG spindle-triggered average always yielded a spindle-like oscillation, confirming the presence of sub-threshold spindles even in the other patients. The small number of patients in our sample precludes a precise explanation of the presence or absence of visible spindles in different patients. A possible explanatory mechanism is implantation location, as both patients with visible spindles had IMEs close to the cerebral midline, which features prominent sleep spindle activity (Andrillon et al., 2011; Fogel and Smith, 2011; Piantoni et al., 2017). Variations in signal quality resulting from the microenvironment of the implanted microelectrodes are another possible explanation for inter-patient differences in IME spindle prominence.

Extremely local spindles were very common not only on ECoG derivations, but also on IME channels. Various intracortical spindle topographies were observed for the micro-domain spindles detected using IME channels. These included the proposed “matrix” spindles in layer I-II and “core” spindles in layer IV (Piantoni et al., 2016) **(Figure 11)**, although these topographies were not particularly common and various other topographies were also seen. These results indicate that the small-scale micro-domain spindle events can be generated by diverse thalamocortical and cortico-cortical networks with various laminar innervation characteristics. In line with previous results (Andrillon et al., 2011; Nir et al., 2011), large-amplitude spindles were detected on more channels **(Figure 3, Panel F-G, Table 2)**, showing that both the extent and amplitude of these oscillations reflect the degree to which their generators are synchronized.

**Figure 11.**
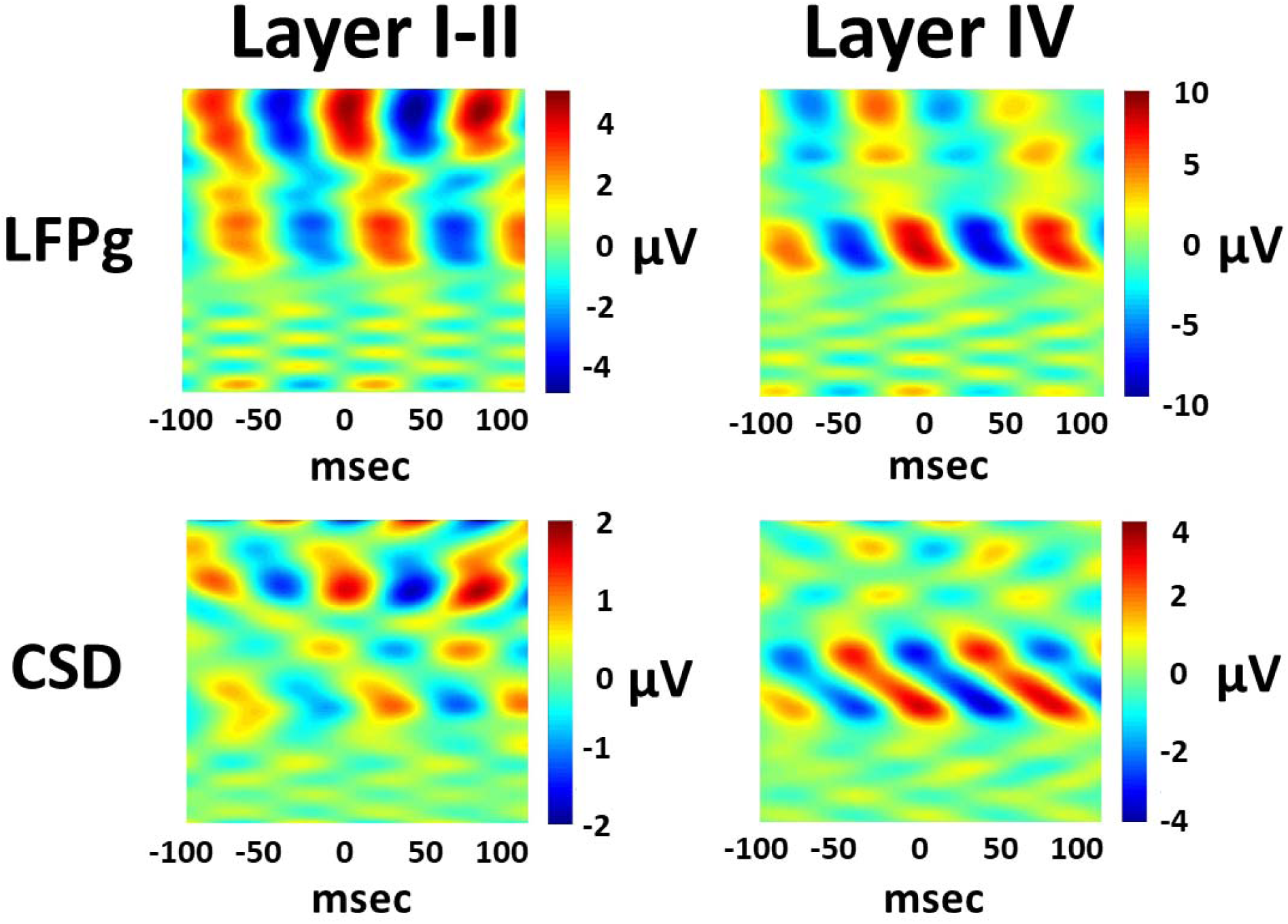
The LFPg and CSD laminar profile of a subset of IME spindles exclusively detected in layer I and II (left panels) and layer IV (right panels) in Patient 2. Data is shown for slow spindles due to the larger number of available events. μV values refer to μV/150 μm in case of LFPg and μV/150 μm^2^ in case of CSD.

The co-occurrence of spindles on IME and ECoG is not symmetrical: macro-domain ECoG spindles are usually (∼90%) accompanied with a spindle in at least one cortical layer, but a relatively large proportion (∼50%) of micro-domain IME spindles did not co-occur with ECoG spindles (Figure 4). This dichotomy is reminiscent of the observation that spindle oscillations in electrophysiological channels with larger spatial receptive fields (EEG, especially monopolar EEG) usually are also seen in channels with smaller receptive fields (MEG, especially gradiometers) but the reverse is not true (Dehghani et al., 2010; Dehghani et al., 2011): that is, a large degree of sleep spindles are extremely local events and monopolar EEG derivations only detect their largest, most synchronized instances. IME data are essentially bipolar EEG recordings from inside the cerebral cortex, with each channel having an extremely small receptive field measurable in hundreds of micrometers at most (Ulbert et al., 2001). Therefore, IME-exclusive spindles likely represent the extreme left tail of the distribution of the spatial extent of sleep spindles, encompassing the most local instances. At this level of resolution, sleep spindles are present with highly variable laminar topographies, but their amplitude is usually higher in the superficial layers. Density, on the other hand, is relatively even across cortical layers - it must be noted, however, that the IAM detection method is especially well adapted to detecting low-amplitude spindles as long as individual frequency ranges are correctly provided using channels with more prominent spindles. Therefore, other detection methods might have detected lower spindle densities on deeper channels.

Our analysis of the laminar profile of the IME signal triggered to macro-domain ECoG sleep spindles revealed large between-subjects differences in absolute voltage and the layer-wise distribution of amplitude, but within the same subject the profile of slow/fast and local/global spindles was very similar **(Figure 6-8).** On average both LFPg and CSD magnitude were maximal in Layer I-IV, and markedly reduced in Layer V-VI, while within-subject differences in the cortical profile generally referred to a different superficial layer in which the magnitude was maximal. Global spindles and to a lesser extent slow spindles were also characterized by generally higher signal magnitude **(Figure 6).** We repeated our analyses of the laminar profile using z-transformed magnitudes across layers in order to correct for the large between-subject differences in magnitude. This method is not able to detect the main effect of between-subject factors on the dependent variable, since all spindles by definition have a mean magnitude of 0 with a variance of 1 across layers, but it is sensitive to the interactions between the within-subject variable Layer and the between-subject factors, that is, spindle type effects on the laminar profile. However, the results were very similar: although significant, the effect size of Layer*Spindle type and Layer*Globality interactions was very small. The absence of a substantial interaction effect of spindle globality and spindle type on the magnitude is further evidenced by the fact that LFPg and CSD magnitudes across layers were highly correlated within spindles **(Figure 8).** The lack of clustering on the scatterplot of spindle magnitudes in different layers also indicates that not only is the laminar profile of slow, fast, global and local spindles similar, but that ECoG spindles are generally not the result of two or more IME spindle subtypes, each characterized by a different laminar profile or even abrupt differences in magnitude. Rather, it appears that spindle amplitude, which is at least partially a function of globality (0.13< R^2^<0.34, **Figure 3, Panel F-G)** has a smooth distribution from low to high values which involves each of the cortical layers in a similar manner. In other words, spindle globality appears to be the strongest modulator of spindle amplitude, affecting amplitude in each layer similarly. Note that the R^2^ values we report for the association between spindle amplitude and spindle globality are lower bounds of the real value due to the limited cerebral mapping and resolution of the implanted electroencephalographic sensors: some spindles may have had a large amplitude because their spatial extent was large in an area only partially mapped and thus underestimated by either ECoG or IME.

In sum, the cortical profile of macro-domain ECoG spindles was usually characterized by widespread and uniform spindle-frequency activity mainly concentrated in the superficial layers, indicating that in macro-domain spindles visible on the ECoG cortical recruitment is more similar. The widespread involvement of layer I-IV suggests the activity of widespread thalamocortical and possible cortico-cortical circuits with relatively even contribution to each spindle. The strong correlation and high loading on a single factor of LFPg and CSD amplitude in various layers during ECoG spindles indicates that the average laminar profile is representative for most cases and not a summation of heterogeneous individual profiles. Furthermore, we found no evidence that the laminar profile of the sleep spindle is associated with either spindle type (slow/fast) or spindle globality (local/global). Contrary to previous hypotheses (Piantoni et al., 2016), the laminar activity profile defined by either LFPg or CSD was very similar for all ECoG spindle types within the same patient (Figure 7). The similar laminar profile of slow and fast spindles is also in line with recent findings showing their similar coupling to thalamic downstates (González et al., 2018), suggesting their functional similarity. It must be noted that our application PCA to IME signal magnitude during ECoG spindles is not directly comparable to a previously used method (Hagler et al., 2018) applying it to the sigma-range IME signal itself. Our method specifically investigated the similarity of the IME signal during ECoG spindles, while the other method estimated the similarity of spindle-frequency IME activity. Taking advantage of the high sampling rate of IME recordings and the presence of visible single-cell discharges on several IME channels, we performed an analysis of SUA and MUA across cortical layers during ECoG spindles. Similar analyses have been previously performed in several other studies (Contreras et al., 1997; Hartwich et al., 2009; Andrillon et al., 2011; Peyrache et al., 2011; Gardner et al., 2013; Sela et al., 2016), with variable results, possibly owing to their methodological differences, most prominently species of subjects (rats, cats or humans), electrophysiological recording methods (tetrodes, extracellular recordings or depth EEG with microwires) and the use of various EEG references, the latter of which makes the interpretation of negative and positive phases particularly difficult. Our results are rather in agreement with the only available human study (Andrillon et al., 2011): we found that SUA was strongest during sleep spindle troughs and the preferred phase of single-unit discharges was during the positive-negative transition of sleep spindles. MUA was also heavily entrained by sleep spindle activity, and exhibited an antiphase correlation pattern with local field potentials across most cortical layers. Our results indicate that it is the derivative, rather than the amplitude of the LFPg signal which is the function of local cell firing, indexed by both SUA and MUA: LFPg phase transitions rather than minima and maxima correspond to SUA/MUA minima and maxima. The imperfect correlation of the two signals indicate that LFPg is a relatively weak function of local neuronal activity, and may rely on extracellular currents and volume conduction from more distant tissue in addition to the action potentials indexed by SUA and MUA. Notably, spindle type, layer and patient differences in the preferred SUA phase angles and the LFPg-MUA relationship indicates that the relative contribution of local neuronal events to the local field potential may vary not only across oscillation types, cortical layers patients, but possibly also over the course of spindles. This observation may have implications about the heterogeneous generating mechanisms of these oscillations. However, due to the lack of consistent effects across the two patients as well as the imperfect statistical power of our study this issue requires further research. It is of note that this study is the first to utilize invasive recordings together with the IAM method. The strong entrainment of MUA/SUA indicators of neuronal activity by IAM spindles is evidence for the reliability of this method.

In sum, our study provides evidence that 1) micro-domain sleep spindles occur with a highly variable spatial extent within the cerebral cortex, 2) micro-domain sleep spindles may occur within any cortical layer, but their layer of occurrence is not systematically different across spindle types 3) the laminar profile of various macro-domain spindle types is also similar and 4) SUA and MUA dynamics are, however, affected by spindle types. These results confirm that while sleep spindles indeed occur with a widespread laminar topography at least at the micro-domain level (Hagler et al., 2018), suggesting the involvement of various thalamocortical and cortico-cortical networks, this network of origin is similar for all spindle types regardless of frequency or spatial extent. Local neuronal dynamics, indexed by SUA and MUA, are however not identical across spindle types (but also layers, patients and individual spindles), rendering them more promising indicators of the functional differences of spindle subtypes.

Our study has a number of shortcomings which may affect the interpretability of our results. First, as it is usual in human invasive electroencephalographic studies, the number of patients used for analyses were low (N=4). Notably, strong inter-patient differences in the laminar profile of sleep spindles were seen, possibly arising from a combination of variable implantation topography and pathological differences in the cerebral cortex. However, the superficial amplitude maximum of sleep spindles and the similarity of the LFPg and CSD laminar profile of various spindle types was a consistent finding across patients. Furthermore, the laminar profile and ECoG-IME occurrence and co-occurrence patterns of sleep spindles were very similar in the two patients (Patient 1 and Patient 2) with visually identifiable IME spindles.

## Funding sources

This work was supported by the following grants:

NAP: 2017-1.2.1-NKP-2017-00002 (NKFIH: National Research, Development and Innovation Office, Hungary)

RO1: 2007P000165/PHS (NIH: National Institute of Health, USA)

Péter P. Ujma was supported by the ÚNKP-17-4 New National Excellence Program of the Ministry of Human Capacities.

The authors declare no conflict of interest.

## Supplementary material

**Supplementary table 1.** ANOVA results for the effects of Patient, Layer, Spindle type and Globality on LFPg signal magnitude, using nominal values (left) and z-transforms across layers (right). Note that between-subject effects could not be estimated using z-transforms because all spindles had a mean magnitude of 0 and a standard deviation of 1 across layers.

| Nominal values | df | F | p | Partial eta-squared | Z-scores | df | F | p | Partial eta-squared |
| --- | --- | --- | --- | --- | --- | --- | --- | --- | --- |
| Intercept | 1 | 4,062.8818 | <0.0001 | 0.5788 | Intercept | 1 |  |  |  |
| {1}Patient | 3 | 3,028.1641 | <0.0001 | 0.7544 | {1}Patient | 3 |  |  |  |
| {2}Global | 1 | 66.1237 | <0.0001 | 0.0219 | {2}Global | 1 |  |  |  |
| {3}Spindle type | 1 | 9.1245 | 0.0025 | 0.0031 | {3}Spindle type | 1 |  |  |  |
| Patient*Global | 3 | 55.0998 | <0.0001 | 0.0529 | Patient*Global | 3 |  |  |  |
| Patient*Spindle type | 3 | 10.8747 | <0.0001 | 0.0109 | Patient*Spindle type | 3 |  |  |  |
| Global*Spindle type | 1 | 1.0181 | 0.3131 | 0.0003 | Global*Spindle type | 1 |  |  |  |
| Patient*Global*Spindle type | 3 | 2.2699 | 0.0785 | 0.0023 | Patient*Global*Spindle type | 3 |  |  |  |
| Error | 2957 |  |  |  | Error | 2957 |  |  |  |
| {4}Layer | 4 | 573.2055 | <0.0001 | 0.1624 | {4}LAYER | 4 | 1,739.5778 | <0.0001 | 0.3704 |
| Layer*Patient | 12 | 697.1830 | <0.0001 | 0.4143 | LAYER*Patient | 12 | 937.6110 | <0.0001 | 0.4875 |
| Layer*Global | 4 | 21.5761 | <0.0001 | 0.0072 | LAYER*Global | 4 | 0.7914 | 0.5306 | 0.0003 |
| Layer*Spindle type | 4 | 3.7308 | <0.0001 | 0.0013 | LAYER*Spindle type | 4 | 2.9564 | 0.0187 | 0.0010 |
| Layer*Patient*Global | 12 | 28.7058 | <0.0001 | 0.0283 | LAYER*Patient*Global | 12 | 3.2047 | 0.0001 | 0.0032 |
| Layer*Patient*Spindle type | 12 | 5.9011 | <0.0001 | 0.0060 | LAYER*Patient*Spindle type | 12 | 3.0284 | 0.0003 | 0.0031 |
| Layer*Global*Spindle type | 4 | 1.7792 | 0.1299 | 0.0006 | LAYER*Global*Spindle type | 4 | 1.5241 | 0.1921 | 0.0005 |
| 4*1*2*3 | 12 | 1.4512 | 0.1349 | 0.0015 | 4*1*2*3 | 12 | 1.4695 | 0.1274 | 0.0015 |
| Error | 11828 |  |  |  | Error | 11828 |  |  |  |

**Supplementary table 2.** ANOVA results for the effects of Patient, Layer, Spindle type and Globality on CSD signal magnitude, using nominal values (left) and z-transforms across layers (right). Note that between-subject effects could not be estimated using z-transforms because all spindles had a mean magnitude of 0 and a standard deviation of 1 across layers.

| Nominal values | df | F | p | Partial eta-squared | Z-scores | df | F | p | Partial eta-squared |
| --- | --- | --- | --- | --- | --- | --- | --- | --- | --- |
| Intercept | 1 | 7633.8165 | <0.0001 | 0.7208 | Intercept | 1 |  |  |  |
| {1}Patient | 3 | 6913.5382 | <0.0001 | 0.8752 | {1}Patient | 3 |  |  |  |
| {2}Global | 1 | 59.1533 | <0.0001 | 0.0196 | {2}Global | 1 |  |  |  |
| {3}Spindle type | 1 | 5.6610 | <0.0001 | 0.0019 | {3}Spindle type | 1 |  |  |  |
| Patient*Global | 3 | 58.3287 | <0.0001 | 0.0559 | Patient*Global | 3 |  |  |  |
| Patient*Spindle type | 3 | 8.8979 | <0.0001 | 0.0089 | Patient*Spindle type | 3 |  |  |  |
| Global*Spindle type | 1 | 4.4534 | 0.0349 | 0.0015 | Global*Spindle type | 1 |  |  |  |
| Patient*Global*Spindle type | 3 | 2.5222 | 0.0561 | 0.0026 | Patient*Global*Spindle type | 3 |  |  |  |
| Error | 2957 |  |  |  | Error | 2957 |  |  |  |
| {4}Layer | 4 | 749.9409 | <0.0001 | 0.2023 | {4} Layer | 4 | 3044.4900 | <0.0001 | 0.5073 |
| Layer *Patient | 12 | 878.9979 | <0.0001 | 0.4714 | Layer *Patient | 12 | 908.9410 | <0.0001 | 0.4798 |
| Layer *Global | 4 | 31.1215 | <0.0001 | 0.0104 | Layer *Global | 4 | 4.5850 | 0.0011 | 0.0015 |
| Layer *Spindle type | 4 | 4.6991 | 0.0009 | 0.0016 | Layer *Spindle type | 4 | 5.6270 | 0.0002 | 0.0019 |
| Layer *Patient*Global | 12 | 33.0743 | <0.0001 | 0.0325 | Layer *Patient*Global | 12 | 7.3643 | <0.0001 | 0.0074 |
| Layer *Patient*Spindle type | 12 | 5.0881 | <0.0001 | 0.0051 | Layer *Patient*Spindle type | 12 | 4.1790 | <0.0001 | 0.0042 |
| Layer *Global*Spindle type | 4 | 3.0812 | 0.0151 | 0.0010 | Layer *Global*Spindle type | 4 | 0.8312 | 0.5051 | 0.0003 |
| 4*1*2*3 | 12 | 2.1036 | 0.0138 | 0.0021 | 4*1*2*3 | 12 | 0.6591 | 0.7921 | 0.0007 |
| Error | 11828 |  |  |  | Error | 11828 |  |  |  |

**Supplementary figure 1.**
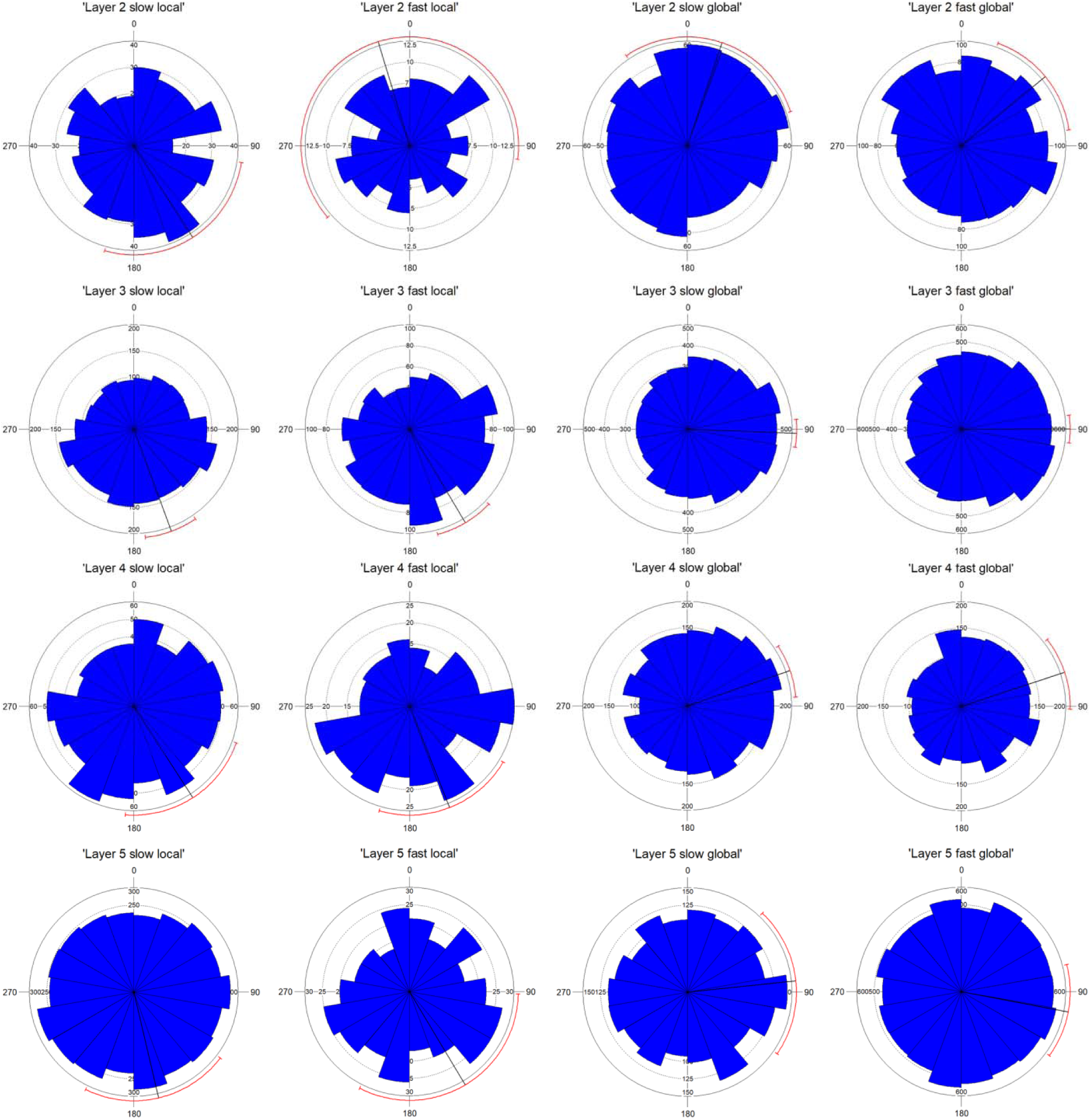
The circular mean of same-channel LFPg phases concomitant to SUA during spindles (±500 msec relative to ECoG spindle events) in Patient 1 by layer and spindle type. All SUA events during spindles of the same type were pooled from all cells and all channels within the same layer. Error bars indicate 95% confidence intervals.

**Supplementary figure 2.**
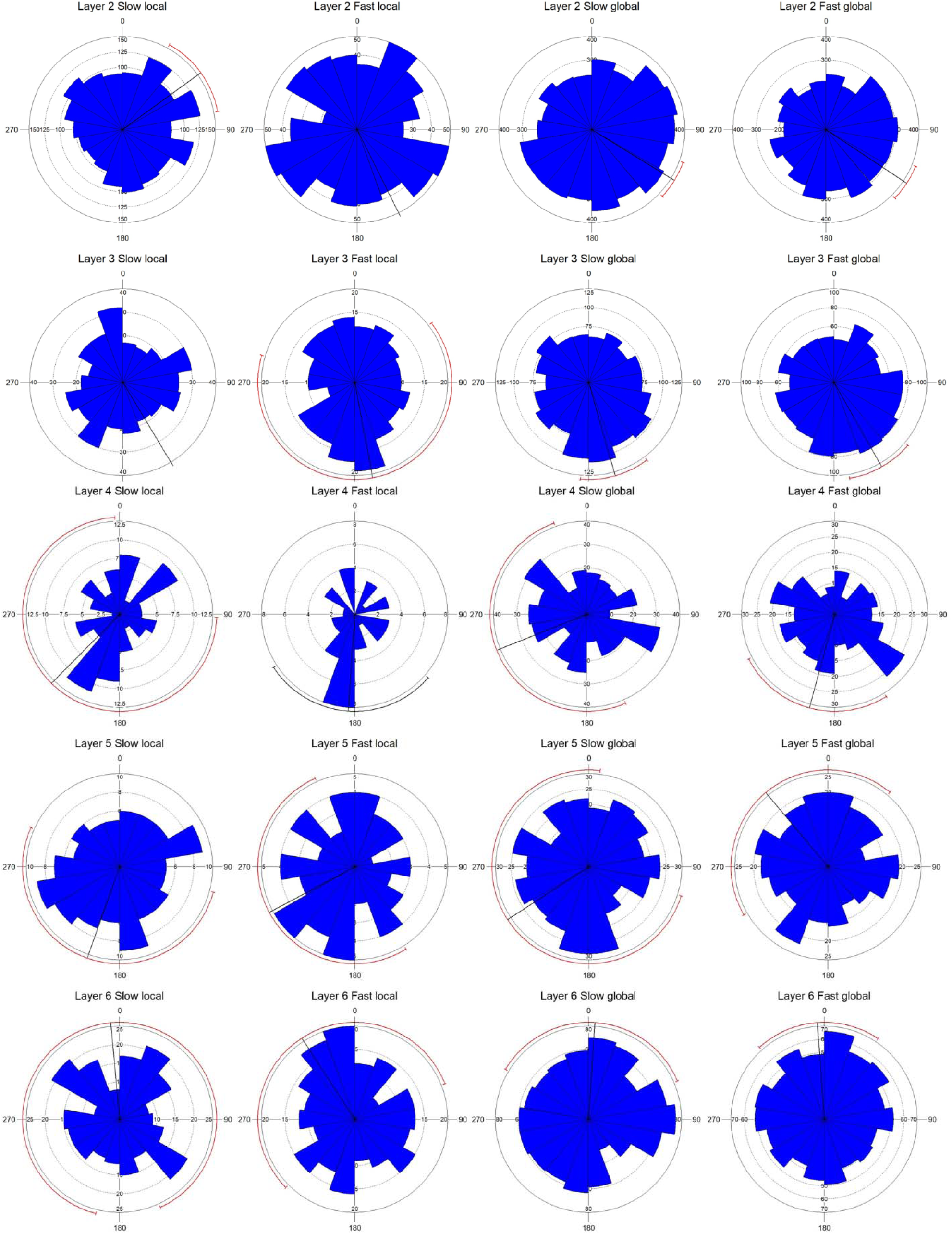
The circular mean of same-channel LFPg phases concomitant to SUA during spindles (±500 msec relative to ECoG spindle events) in Patient 2 by layer and spindle type. All SUA events during spindles of the same type were pooled from all cells and all channels within the same layer. Error bars indicate 95% confidence intervals.

